# Cross-species orthology detection of long non-coding RNAs (lncRNA) through 13 species using genomic and functional annotations

**DOI:** 10.1101/2024.10.03.616473

**Authors:** Fabien Degalez, Coralie Allain, Laetitia Lagoutte, Frédéric Lecerf, Sandrine Lagarrigue

**Author notes:** In the order of authorship, to whom correspondence should be addressed. Tel: NA; /.

## Abstract

Long non-coding RNAs (lncRNAs), defined by a length of over 200 nucleotides and limited protein-coding potential, have emerged as key regulators of gene expression. However, their evolutionary conservation and functional roles remain largely unexplored. Comparative genomics, particularly through sequence conservation analysis, offers a promising approach to infer lncRNA functions. Traditional methods focusing on protein-coding genes (PCGs) fall short due to the rapid evolutionary divergence of lncRNA sequences. To address this, a workflow combining syntenic methods and motif analysis via the Mercator-Pecan genome alignment was developed and applied across 13 vertebrate species, from zebrafish to various amniotes and birds. Further analyses to infer functionality revealed co-expression patterns through 17 shared tissues between human and chicken but also functional short-motif enrichment across the 13 species using the LncLOOM tool, exemplified by the human OTX2-AS1 and its counterparts in other species. The study expanded the catalog of conserved lncRNAs, providing insights into their evolutionary conservation and information related to potential functions. The workflow presented serves as a robust tool for investigating lncRNA conservation across species, supporting future research in molecular biology to elucidate the roles of these enigmatic transcripts.

## INTRODUCTION

Over the last decades, tens of thousands of genes encoding long noncoding RNAs (lncRNAs), characterized by a minimum length of 200 nucleotides and low protein-coding potential, have been identified in genomes [1, 2]. These transcripts have emerged as crucial players in the regulation of gene expression and are involved in multiple biological processes, even if the functions of most lncRNAs remain unknown [3].

Comparative genomics using sequence conservation analysis is an efficient way of inferring function, as is classically done for protein-coding genes (PCGs). However, lncRNA sequences evolve faster compared to most PCGs, which limits their conservation between species, even when synteny is evident. Approximately 70% of lncRNAs have no sequence ortholog in species that have diverged for over 50 million years meaning, for example, that around 100 lncRNAs seems conserved between mammals and fish [4]. As a result, no lncRNA ortholog are reported in reference databases such as Ensembl, in contrast to PCGs. Concerning sequence conservation, different studies report that only a few units of the primary sequence would be kept in order to maintain functional aspects. More specifically, sequence conservation seems to occur in blocks of 5 to 30 nucleotides, known as k-mers [4–6]. For instance, the lncRNA OIP5-AS in human, also referred as Cyrano in fish, contains a 300-nucleotides conserved segment in tetrapod, which contains a deeply conserved 26-nucleotides motifs that binds miR-7 and facilitates the degradation of the microRNA [7]. Another example is CHASERR, recently shown to share two highly conserved 7-nucleotides motifs in the last exon, which have been shown to bind specific proteins [8]. In this context, two primary approaches have been employed for comparative genomic analyses of lncRNAs. These firsts involve analyzing their positional conservation (synteny) within the genome of each species by identifying positionally conserved neighboring PCGs. Some studies have identified positionally conserved lncRNAs within more or less phylogenetically distant species [4, 9], including farm animal species and chicken that diverged from mammals 300 million years ago [10–12]. The second approach consists in searching for conserved short motifs between species, which requires alternatives to conventional sequence algorithms used for PCGs that search for significantly long and highly conserved sequences. Different alternatives are then reported in the literature. SEEKR uses the abundance patterns of short k-mers to assess the functional similarity across sequences. These k-mers could align with certain protein binding sites, and studies have demonstrated the efficacy of SEEKR in clustering disparate lncRNAs with presumably correlated roles within a single species. Another recently developed solution is LncLOOM (LncRNA Linear Order cOnserved Motifs) [8]. This tool is based on the identification of combinations of short motifs (6–12 nucleotides) found in the same order in putatively homologous sequences from different species. This algorithmic framework, by construction, focuses on a user-defined lncRNA and can efficiently compare dozens of putative orthologous sequences from different species for the lncRNA of interest. In addition, LncLOOM can annotate conserved motifs based on miRNA binding site using TargetScan [13]. Finally, the lncHOME computational pipeline identifies, at the genome scale, lncRNAs with conserved genomic positions and short motifs of RNA-binding proteins (RBPs), initially defined from experimental studies and databases [14]. While LncLOOM cannot carry out analyses at the scale of complete genomes, LNChome hypothesizes which functional sequences to be searched for.

As a complement to these tools, we have developed a workflow to identify orthologous lncRNAs on a genome-wide scale, between different user-defined species. Such a workflow combines three methods: two syntenic methods and one method analyzing short motifs using the “Mercator-Pecan” genome alignment method [15, 16] used by Ensembl. These approaches have been applied to a range of 13 species, including the zebrafish and 12 amniotes, from *Boreoeutheria* and *Aves*, which diverged around 320 million years ago and which include the main farm and domesticated species (cow, pig, chicken). To go further, orthologous lncRNAs can be deeply analyzed using LncLOOM, as illustrated by the example of the human lncRNA OTX2-AS1, or by applying two additional functional analyses based on gene expression, illustrated in this paper with orthologous lncRNAs between human and chicken. The first one consists in analyzing co-expression across-tissues of orthologous lncRNA using a set of common tissues in the species studied, here 17 tissues for human and chicken. The second one consists in evaluating the intra-tissue enrichment between the chicken and the human of co-expressed PCGs with each putative orthologous lncRNAs. Such analyses can reveal useful functional information, to support further molecular biology experiments aimed at deciphering lncRNA function.

Together, our study substantially expands the known repertoire of conserved lncRNAs across 13 vertebrates, offering insights into their evolutionary conservation and function and provides a workflow to investigate lncRNA conservation in other species.

## RESULTS

### Genome annotation variability and orthology of PCG in 13 vertebrates

A total of 13 species were used for the analysis (Figure 1A). This set includes the zebrafish and 12 amniotes, which diverged around 429 million years ago. Within the amniote group, species from *Boreoeutheria* and *Aves*, which diverged around 320 million years ago, were selected. For the *Boreoeutheria* group, farm and domestic species, such as the horse, cow, pig, goat, and dog were considered, along with reference species such as the human and mouse. In the *Aves* group, the chicken was included due to its importance in research and agriculture, along with the duck and turkey. Additionally, the zebra finch was added as a well-annotated representative of the *Aves*. Finally, as the cow and the goat which have only few numbers of annotated lncRNAs (1,480 and 2,705 respectively), the ostrich, with 1,034 lncRNAs identified, was added to the *Aves* group (Figure 1B). More generally, an overview of genome annotations revealed substantial variation in the number and characteristics of annotated genes across these 13 species. As shown in Figure 1B, the total count of long non coding RNA (lncRNA) loci ranged from 1,034 for the turkey to 18,859 in the human and to 44,428 in *the chicken* for which a custom gene-enriched annotation was used in addition to the Ensembl annotation. Note that the median number of lncRNA across species is 6,479. For protein coding genes (PCG), the number varies from 14,297 in *the ostrich* to 25,432 in the zebrafish with a median of 21,343 genes. Despite the disparities in lncRNA counts, the annotated lncRNAs displayed similar structural properties across most species. The mean proportion of monoexonic (single-exon) lncRNA loci hovered around 13%, with the exception of *the chicken* enriched version at 38% (Sup. Figure 1). Similarly, the average number of annotated lncRNA isoforms (transcripts per locus) was fewer than 2 for the majority of species, with slightly higher values observed in Homo sapiens (> 5), Mus musculus, and Equus caballus (> 3) (Sup. Figure 1) due to difference in annotation effort.

**Figure 1.**
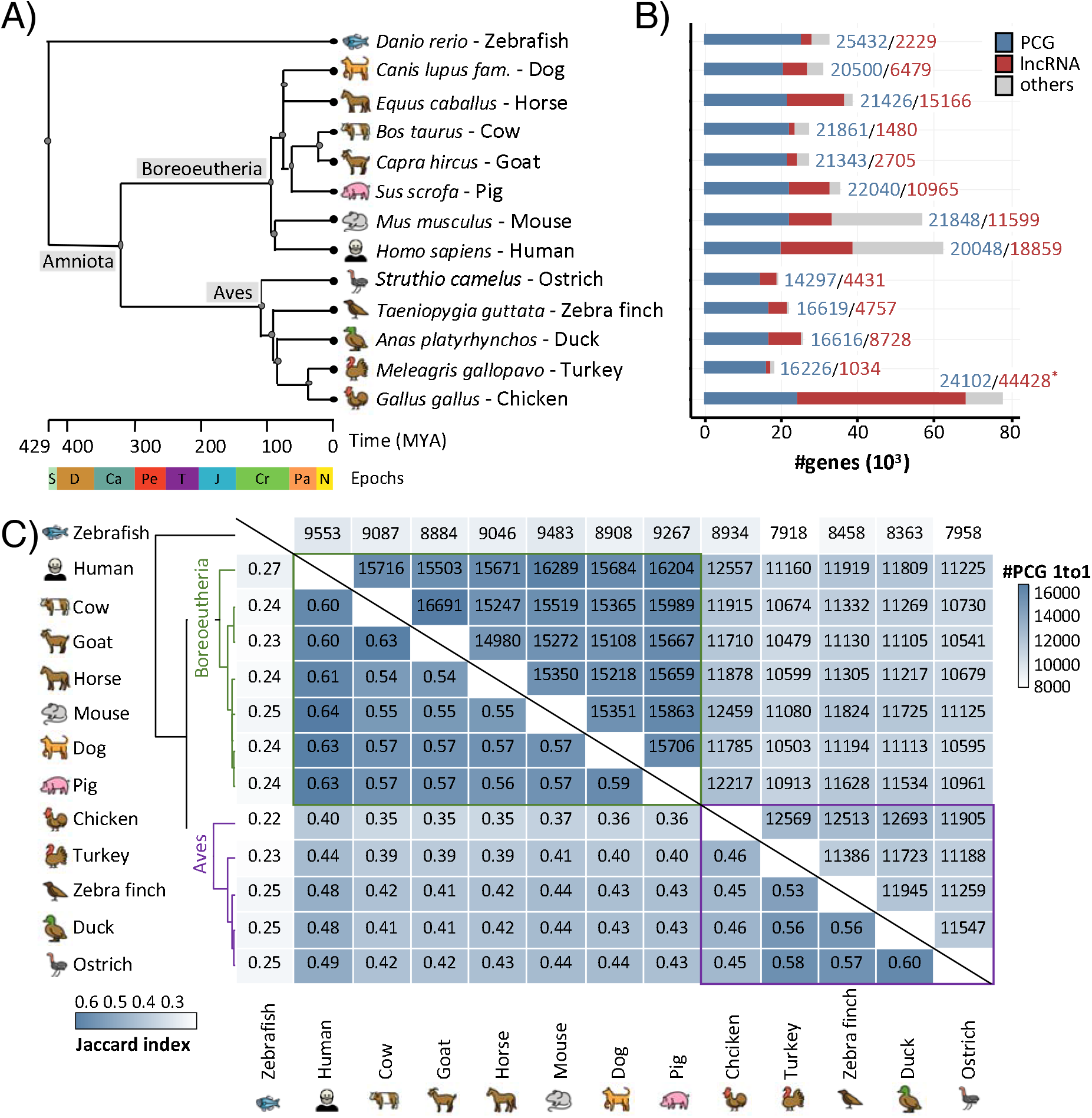
Genome annotation characteristics of the 13 species. (a) Phylogenetic tree based on the molecular time estimates in Timetree [49]. S: Silurian, D: Devonian, Ca: Carboniferous, Pe: Permian, T: Triassic, J: Jurassic, Cr: Cretaceaous, Pa: Paelogene, N:Neogene. (b) Number of PCGs (blue), lncRNAs (red) and other (grey) genes identified in each genome annotation. *: custom gene-enriched annotation for the chicken. (c) Clustered heatmap based on pairwise Jaccard index (lower triangle) for one-to-one orthologous PCGs. The upper triangle indicates the raw number of one-to-one orthologous PCGs.

To establish a baseline for comparing lncRNA conservation, the orthologous relation for PCGs across the 13 species was analysed. Pairwise comparisons revealed that the Jaccard index (see Mat and Meth) of one-to-one orthologous PCGs ranged from 0.22 between *the chicken* and zebrafish (8,934 one-to-one PCGs) to 0.64 between the human and the mouse (16,289 one-to-one PCGs) with a mean of 0.45 (Figure 1C). These proportions aligned with the expected phylogenetic relationships with more closely related species exhibiting higher levels of one-to-one orthology conservation. Note that the Jaccard index averages 0.58 and 0.52 for the *Boreoeutheria* and *Aves* groups respectively.

### Identification of orthologous lncRNAs

#### By “orthology-by-synteny” using the two orthologous flanking PCGs (Figure 2A– 2D)

After considering the baseline for PCG orthology conservation, the orthology relationships of lncRNA genes across the 13 species was then investigated using their flanking one-to-one orthologous PCGs in upstream and downstream. This “orthology-by-synteny” method is based on the hypothesis that lncRNAs located between well-conserved orthologous PCGs may themselves be orthologous, despite potentially rapid sequence evolution. For the pairwise comparison of the 13 species, the percentage of lncRNA flanked by two orthologues PCGs was highly variable going from 13% to 76% for the “zebrafish- turkey” and “mouse-human” pairs respectively, and with a median at 51% (see Sup. Table 1). Note that the number of lncRNA surrounded by only 1 orthologous PCG is around 35%. In terms of lncRNA orthologous relationships, the number of lncRNAs that can be associated with one or more lncRNAs in the other species ranges from 0.22% (turkey – zebrafish; n = 5 – 10) to 40% (human – mouse; n = 4693 – 5829) with a median of 9% (see Table 1 and Sup. Table 2). Note that the “one-to-one” case represent around 2% of the total lncRNA identified as conserved between pairwise species.

**Figure 2.**
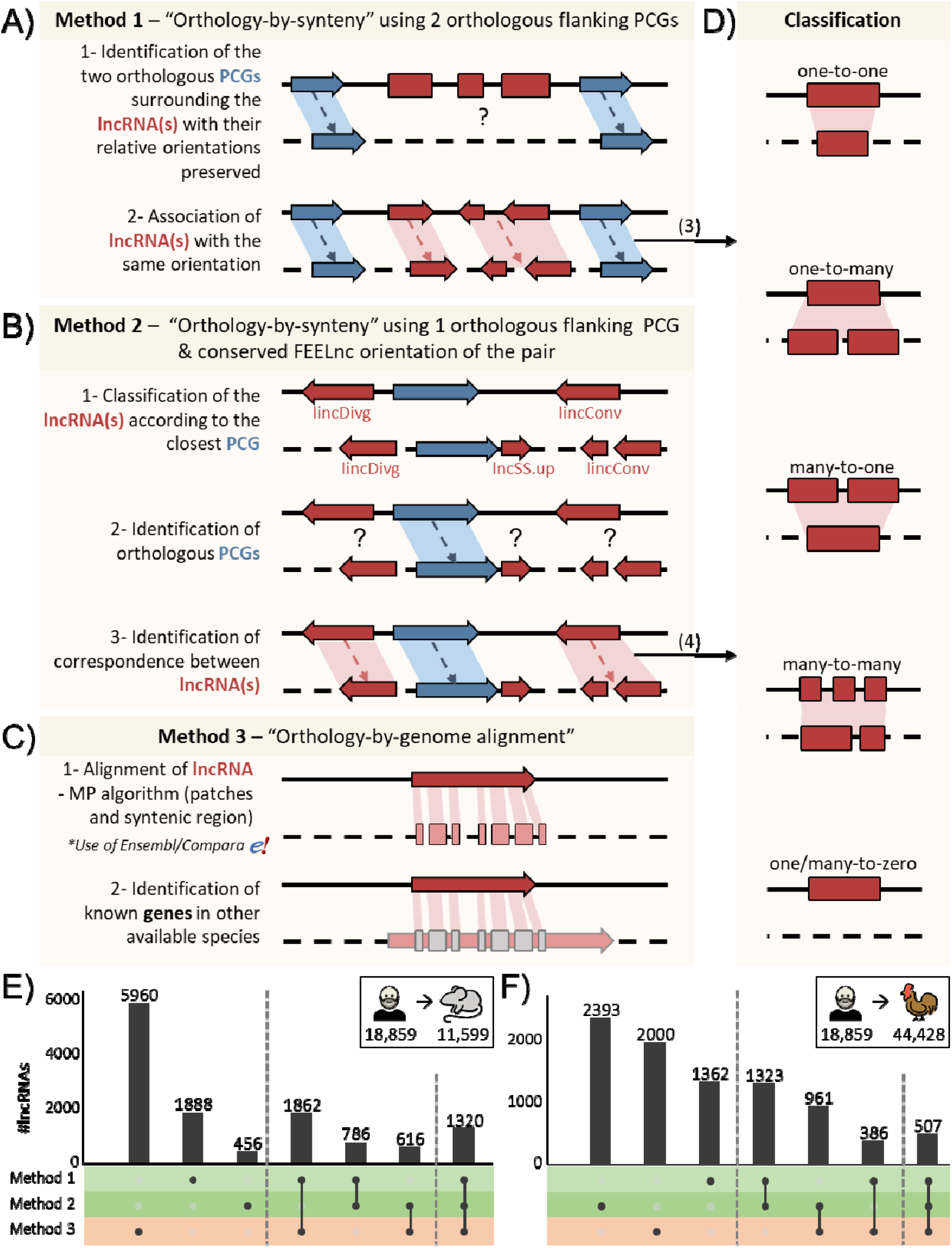
Overview of the methods used. Orthology inference of lncRNAs by examining the synteny using (a) two flanking one-to-one orthologous PCGs or (b) one orthologous lncRNA and the conservation of the FEELnc configuration with the nearest one-to-one orthologous PCG. Configuration classes considered in (b) are detailed in Sup. Figure 2. Orthology by genome alignment using (c) the lncRNA patch matches produced by the Mercator-Pecan algorithm from the “63 amniotes” groups established by Ensembl (v111). (d) Classification of putative orthologous lncRNAs used for methods 1 (a) and 2 (b). LncRNAs and PCGs are represented in red and blue respectively. The arrow orientation corresponds to the gene strand. The plain and dotted lines correspond to the source and target species respectively. (e)- (f) Number of lncRNAs detected by the combination of each methods using the human as reference and the mouse and the chicken as target species respectively.

**Table 1.**
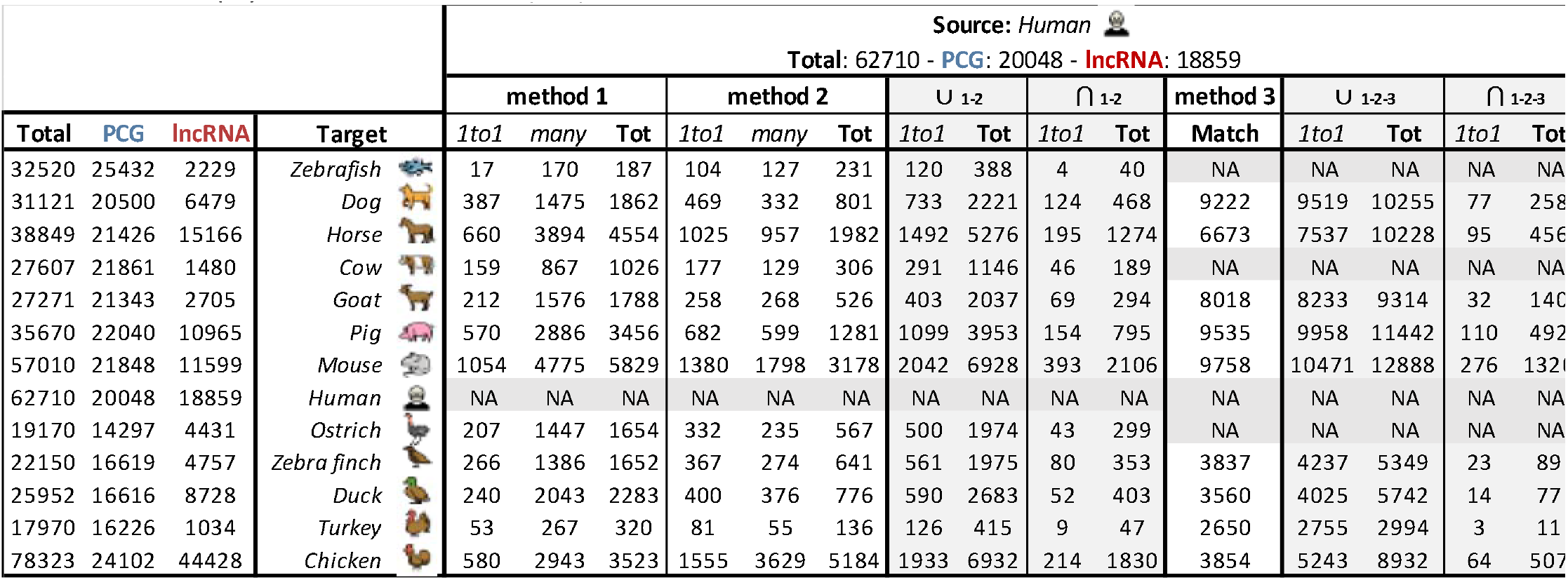
– Number of lncRNA orthologous relations found by each method between the human and the 12 species. For method 1 and method 2, “1to1” refers to lncRNA with a “one-to-one” ortholog and “many” for lncRNA with a “many-to-one”, “one-to-many” or “many-to-many” relation. Union (∪) and intersection (⋂) for method 1 and 2 (1-2) and all methods (1-2-3). NA: Not Applicable.

|  |  |  |  | Source: Human |  |  |  |  |  |  |  |  |  |  |  |  |  |  |
| --- | --- | --- | --- | --- | --- | --- | --- | --- | --- | --- | --- | --- | --- | --- | --- | --- | --- | --- |
|  |  |  |  | Total: 62710 - PCG: 20048 - lncRNA: 18859 |  |  |  |  |  |  |  |  |  |  |  |  |  |  |
|  |  |  |  | method 1 |  |  | method 2 |  |  | U 1-2 |  | ∩ 1-2 |  | method 3 | U 1-2-3 |  | ∩ 1-2-3 |  |
| Total | PCG | lncRNA | Target | 1to1 | many | Tot | 1to1 | many | Tot | 1to1 | Tot | 1to1 | Tot | Match | 1to1 | Tot | 1to1 | Tot |
| 32520 | 25432 | 2229 | Zebrafish | 17 | 170 | 187 | 104 | 127 | 231 | 120 | 388 | 4 | 40 | NA | NA | NA | NA | NA |
| 31121 | 20500 | 6479 | Dog | 387 | 1475 | 1862 | 469 | 332 | 801 | 733 | 2221 | 124 | 468 | 9222 | 9519 | 10255 | 77 | 258 |
| 38849 | 21426 | 15166 | Horse | 660 | 3894 | 4554 | 1025 | 957 | 1982 | 1492 | 5276 | 195 | 1274 | 6673 | 7537 | 10228 | 95 | 456 |
| 27607 | 21861 | 1480 | Cow | 159 | 867 | 1026 | 177 | 129 | 306 | 291 | 1146 | 46 | 189 | NA | NA | NA | NA | NA |
| 27271 | 21343 | 2705 | Goat | 212 | 1576 | 1788 | 258 | 268 | 526 | 403 | 2037 | 69 | 294 | 8018 | 8233 | 9314 | 32 | 140 |
| 35670 | 22040 | 10965 | Pig | 570 | 2886 | 3456 | 682 | 599 | 1281 | 1099 | 3953 | 154 | 795 | 9535 | 9958 | 11442 | 110 | 492 |
| 57010 | 21848 | 11599 | Mouse | 1054 | 4775 | 5829 | 1380 | 1798 | 3178 | 2042 | 6928 | 393 | 2106 | 9758 | 10471 | 12888 | 276 | 1321 |
| 62710 | 20048 | 18859 | Human | NA | NA | NA | NA | NA | NA | NA | NA | NA | NA | NA | NA | NA | NA | NA |
| 19170 | 14297 | 4431 | Ostrich | 207 | 1447 | 1654 | 332 | 235 | 567 | 500 | 1974 | 43 | 299 | NA | NA | NA | NA | NA |
| 22150 | 16619 | 4757 | Zebra finch | 266 | 1386 | 1652 | 367 | 274 | 641 | 561 | 1975 | 80 | 353 | 3837 | 4237 | 5349 | 23 | 89 |
| 25952 | 16616 | 8728 | Duck | 240 | 2043 | 2283 | 400 | 376 | 776 | 590 | 2683 | 52 | 403 | 3560 | 4025 | 5742 | 14 | 77 |
| 17970 | 16226 | 1034 | Turkey | 53 | 267 | 320 | 81 | 55 | 136 | 126 | 415 | 9 | 47 | 2650 | 2755 | 2994 | 3 | 11 |
| 78323 | 24102 | 44428 | Chicken | 580 | 2943 | 3523 | 1555 | 3629 | 5184 | 1933 | 6932 | 214 | 1830 | 3854 | 5243 | 8932 | 64 | 507 |

#### ***-*** By “orthology-by-synteny” using one orthologous flanking PCGs and the conserved FEELnc orientation of the lncRNA-PCG pair (Figure 2B– 2D)

To complement the previous approach which required in the annotation file the presence of the two flanking orthologous PCGs, we looked at the lncRNAs according to the presence of only one flanking orthologous PCG. However, this filter was consolidated by adding a second criteria based on the FEELnc configuration consisting in analysing the orientation and distance of the lncRNA-PCG pair (see Sup. Figure 2). Among the lncRNA classified and due to difference in annotation, note that a disparity is observed between the “genic” and “intergenic” classes and also between the four constitutive subclasses: convergent, divergent, same strand upstream and downstream (see Sup. Figure 3). The percentage of lncRNA classified by FEELnc ranged from 62% (n=1,676) in the goat to 98% (n=43,617) in the chicken, with a median of around 82% (see Sup. Table 3). In terms of lncRNA orthologous relationships, the number of lncRNAs that can be associated with one or more lncRNAs in the other species ranges from 0.45% (zebrafish – turkey; n = 10 – 11) to 27% (human – chicken; n = 5184 – 9576) with a median of 4% (see Table 1). Note that the “one-to-one” case represent around 3% of the total lncRNA identified as conserved between pairwise species.

#### ***-*** By “orthology-by-genome alignment” using Ensembl/Compara and the Mercator PECAN alignment of amniotes group (Figure 2C)

Given the low sequence conservation of lncRNAs, conventional alignment methods could not be applied, especially in an automated way. One approach consisted in observing conserved patterns in the sequence of a hypothetical common ancestor obtained via multiple alignment of a group of 63 amniotes (Compara/Ensembl). As specified in the “Material and Methods”, note that this group included 10 of the 13 species analyzed here: the cow, the ostrich and the zebrafish are excluded. Globally, the percentage of lncRNAs with an equivalence in another species ranged from 7% (lncRNAs horse → genome turkey; n = 1091) to 54% (lncRNAs dog → genome pig; n = 3485), with a median of 23% (see Table 1). This number is highly variable according to the phylogenetic distance separating the species concerned, and the *Boreoeutheria* and *Aves* groups thus stand out clearly. More precisely, the median percentage of lncRNAs aligning within these groups is 43% and 34% respectively, while it is 17% between. It should be noted that, unlike the other two methods, this third method only depends on the annotation of the query genome, thus reducing the impact genome annotation.

#### Combined use of the three methods between two species (Figure 2 E)

While each method can be applied independently and for each pair of species, the interest also lies in their complementarity. Thus, using at least one of the 3 methods, it was possible to identify 12,888 (68%) lncRNAs as potentially conserved between human and mouse and with 1,320 (7%) lncRNAs detected by all 3 methods (Table 1 and Figure 2E and Sup. Table 2). However, by construction, only the first two methods allow to presume a relationship between one or more lncRNAs in each species, consequently 2,106 (2%) lncRNAs in humans were detected by both methods by synteny, 393 have a "one_to_one" equivalent in each method and 276 aligned on the mouse genome. Interestingly, among these 276 genes, 160 (58%) have no gene name and 90 (33%) are identified as divergent (-DT) or antisense (-AS). Due to a more important phylogenetic divergence, the number of potentially orthologous lncRNAs found between human and the chicken is lower, with 8,932 (47%) lncRNAs found by at least one method and 507 (3%) by all 3 methods applied together (Table 1 and Figure 2F and Sup. Table 2). Similarly, 1,830 (10%) human lncRNAs were detected by both synteny methods, 214 (1%) had a "one_to_one" equivalent by both methods and 64 are observed through the Mercator-Pecan alignment. In this case, 34 (57%) are not named and 26 (41%) are antisense/divergent, similar proportion as thus observed for the mouse. Interestingly, six lncRNAs (ENSG00000229739, ENSG00000235527, ENSG00000285736, ENSG00000234055, ENSG00000256546, ENSG0000025840) are identified as “one_to_one” by the 2 methods and with an alignment between the human, the mouse and the chicken and 3 have a gene name assigned: PDC-AS1; HIPK1-AS1 and LINC02985.

### Cross-species use of the three methods to identify potential lncRNA orthologs: the example of OTX2-AS1

Applying all three methods to all 13 species provides additional information to support potential orthologous lncRNA relationships. As an example, the OTX2-AS1 antisense lncRNA (ENSG00000248550), annotated in the human which was used as a reference, is identified as potentially conserved in several of the species of the study. Indeed, as illustrated in Figure 3A (and Sup. Table 4), this lncRNA was initially identified in 9 of the 12 other species using one or more methods, each providing clues to the orthology relationship. Concerning method 1, OTX2-AS1 is surrounded downstream by OTX2 and upstream by EXOC5, two PCGs conserved by a one-to-one orthology relationship in all other species. Using this method, several potential orthologs are identified in the mouse, the chicken and the pig, and only one in for the duck. It should be noted that method 1 failed to identify any ortholog relationships in dog and zebrafish, as the latter have a non-conserved PCG upstream of EXOC5. According to method 2, the OTX2- AS1 lncRNA is associated with the PCG OTX2 with a relationship of type “lncgDivg” once the relaxed FEELnc nomenclature has been applied. This method then associates this lncRNA with a single ortholog in mouse, chicken, pig, duck, dog and zebrafish. Note also that the associated lncRNA in mice is named Otx2os, supporting the potential orthology relationship. In other species, the concerned lncRNA in never has a gene name according to the HGNC nomenclature. Finally, method 3 (applied to 10 available species) identified alignment zones for the mouse, the chicken, the duck and the dog, but also in species where no other method showed results, such as the goat, turkey and zebra finch. Detailed observation of the Mercator-Pecan multiple alignment (Figure 3B) allows to identify two different alignment patterns corresponding to *Boreoeutheria* and *Aves* respectively. However, even if these patterns show a large number of dissimilarities, some areas appear to be preserved, which might indicate areas of functional importance. Finally, note that for the 3 species for which no method was able to provide any information, *i.e.*, the cow, horse and ostrich, no lncRNA was annotated in the genomic area delimited by OTX2 and EXOC5.

**Figure 3.**
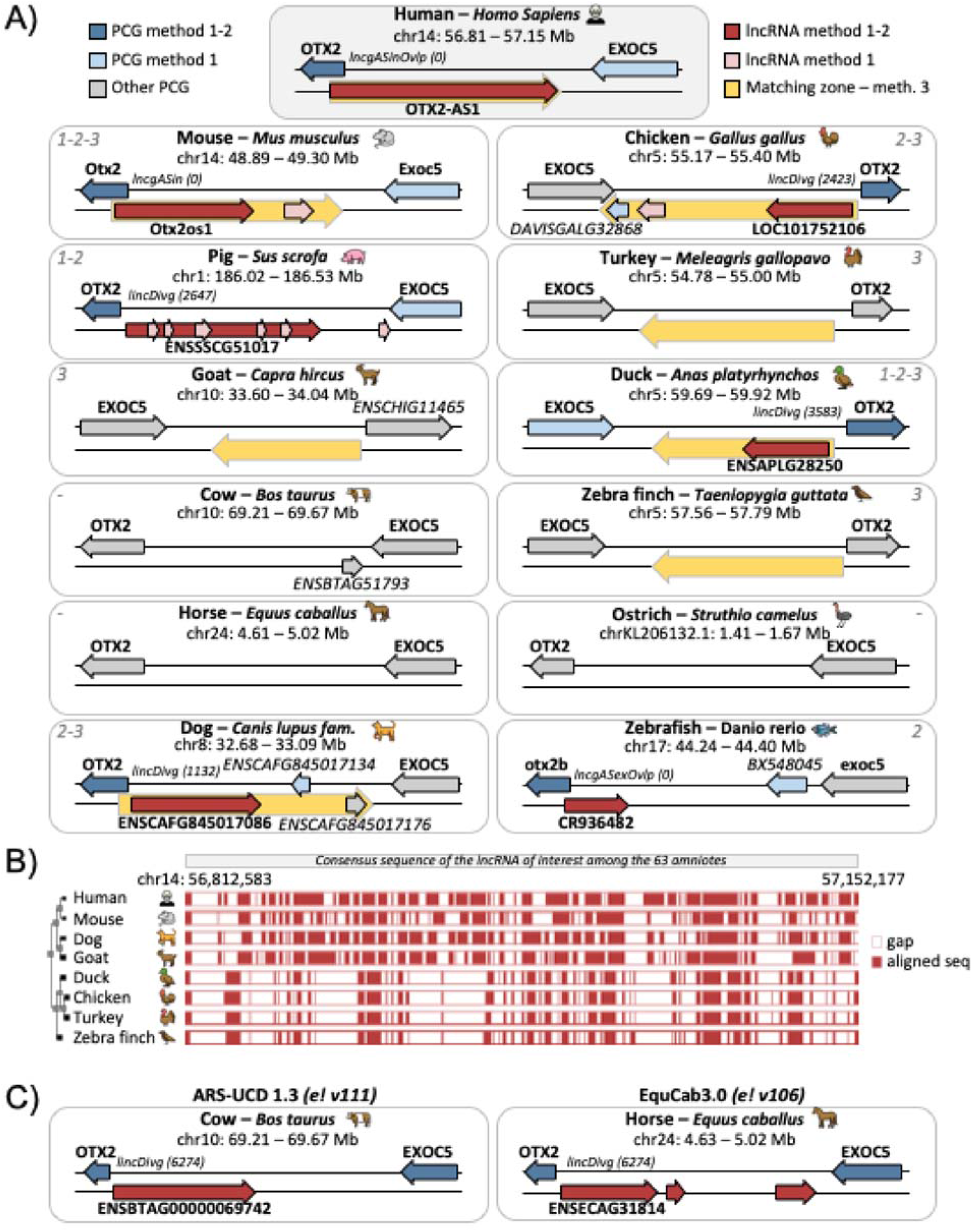
Identification of putative lncRNA orthologs for human lncRNA OTX2-AS1 across 13 species. (a) Genomic regions and results of the three methods considered for research of potential orthologs for OTX2-AS1 (indicated by the grey top numbers). The arrow orientation corresponds to the gene strand. The number associated to the configuration corresponds to the distance between the lncRNA and the PCG. (b) Alignment details generated by the Mercator-Pecan algorithm for available species. The human sequence of OTX2-AS1 was used as input. (c) Genomic regions associated with potential OTX2-AS1 ortholog considered after manual observation. Details are provided in Sup. Table 4.

As shown, the analysis of putative orthology relationships between lncRNAs is annotation-dependent, and manual observations can complement the results of the applied workflow. Thus, for the cow, the release of the new ARS-UCD 1.3 assembly (previously ARS-UCD 1.2) and its annotation according to Ensembl v111 led to the annotation of a single new gene, contained between OTX2 and EXOC5, in “lincDivg” configuration relatively to OTX2 and therefore potentially orthologous to OTX2-AS1 (Figure 3C- left). Similarly, for the horse, while no lncRNA is annotated in the new versions of the annotation, it is in an older version (Ensembl v106, for the same EquCab3.0 assembly) that we find 3 lncRNAs in the study zone, including one classified as “lincDivg” in relation to OTX2 (Figure 3C-right). By supplementing the workflow results with a manual analysis, the number of species with OTX2-AS1 conservation clues rises from 9 to 11 with only the ostrich presenting no information. Please note that, although lncRNAs in the same strand position of a PCG, and downstream in particular, are to be considered more cautiously as they may be poorly modeled 3’UTR regions of a PCG, they are not to be evicted.

### Additional analyses for orthologous lncRNAs to infer functionality (case example: OTX2- AS1 / LRRC3B)

#### Identification of short motifs conservation (k-mers) via LncLOOM (Figure 4A)

Based on the results exposed in the Figure 3, exonic sequences of the OTX2-AS1 human lncRNA and the 8 other putative orthologous lncRNAs were extracted and analyzed through the LncLOOM software. Note that the length of sequences provided was highly variable with a minimum of 549 bp for the dog (1 transcript with 3 exons) to 15,504 pb for the pig (5 transcripts with 2 to 7 exons) (mean: 5,100 bp, sd: 5,911). When using the human sequence as reference, a total of 73 conserved motifs were identified (a restricted motif in a set contained in a larger motif in a sub-set is counted only once) and more than 85% were found only between the human and the pig, the second species after the sequences have been reordered based on their homologies (Sup. Table 5). It should be noted that patterns are searched incrementally; therefore, if a pattern is not found in one species, it is not searched in the most distant species. As a results, the “AAAAGTTG” motif of size 8 is found in all species and is included in two other motifs of size 9 and 75 found in 7 species (excluding zebrafish) and 2 species (human and pig) respectively. Interestingly, using the chicken lncRNA sequence (LOC101752106) as the reference, the “AAAAGTT” motif, of size 7, and included in the previous one, was found among a total of 54 motif with 52% (n=28) shared between the chicken and the duck only. Using the human as reference, 19 of the 73 (26%) conserved motifs are associated to a total of 21 potential miRNA binding sites (detected by TargetScan). Note that if more than 80% of miRNA binding sites are identified only in humans and pigs (depth = 2), miR-320 and miR-1251-5p are identified in the duck (depth = 3) and miR-186-5p also in the chicken (depth = 4). Interestingly, this miR-186-5p is found to be associated to 4 different conserved motifs. Interestingly, this miR-186-5p is found to be associated to 4 different conserved motifs and seems to be associated to the “melanogenesis” pathway and in basic processes of neuronal plasticity affected in neurodevelopmental disorders [17], fact coherent with the expression pattern observed for OTX2-AS1 across various tissues. The enrichment analysis based on miRNA also showed an enrichment in the “retinoblastoma” (n=9; p-val = 2.95e-4) and “brain disease” (n=16; p-val = 7.07e-5) diseases (MNDR - mammal ncRNA–disease repository) category (see Sup. Table 6). Note that OTX2 is involved in the development of the retina, photoreceptors and pineal gland [18] and that OTX2 and OTX2-AS1 transcription is regulated from a shared bidirectional promoter [19].

**Figure 4.**
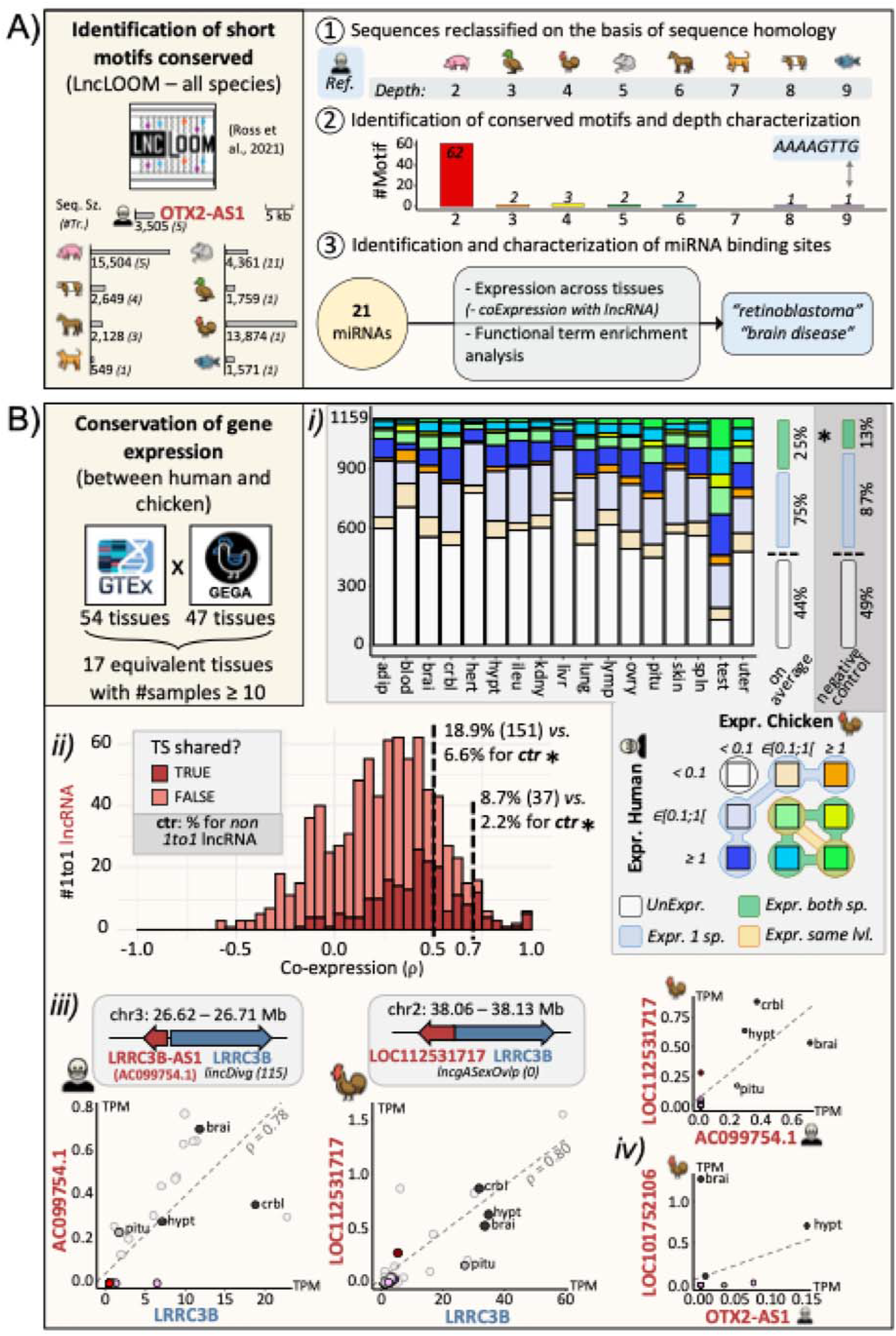
Additional analyses for orthologous lncRNAs to infer functionality. A) Identification of short motif conserved across the 13 species for the OTX2-AS1 lncRNA and its putative orthologs using LncLOOM [8, 44] and enrichment analysis for miRNA through miEAA [38]. B) Conservation of gene expression across 17 tissues between the chicken and the human. *i)* Proportion of the 1,159 putative 1to1 orthologous lncRNAs between the human and the chicken regarding expression categories (TPM < 0.1; ∈ [0.1;1[ or ≥ 1). *ii*) Distribution of lncRNA detected as tissue-specific for at least one common tissue according to the co-expression. *iii)* Expression conservation between the LRRC3B PCG and its antisense lncRNA identified for the human and the chicken. Transparent dots correspond to all the tissues available in the species-specific datasets, while the colored dots correspond to the 17 common tissues identified. iv) Expression conservation between the human lncRNA OTX2-AS1 and its putative ortholog in the chicken. Tissues abbreviations are available in the Sup. Table 7.

#### Conservation of gene expression between the human and chicken (Figure 4B)

As detailed in the Material and Methods part, a total of 17 tissues, considered as equivalent between the human and the chicken has been studied (Sup. Table 7). Concerning the human, 11,971 (88%) lncRNAs and 18,587 (97%) PCGs are considered as expressed in at least one of the tissues (TPM ≥ 0.1). A total of 7,054 lncRNAs are considered as TS (τ ≥ 0.9) with 4,907 (70%) with the testis as the most expressed tissue. Parallelly, for the chicken, 28,384 (66%) lncRNAs and 17,688 (87%) PCGs are expressed and 20,163 lncRNAs are considered as TS always with the testis as the tissue supporting the more TS genes with a total of 7,891 lncRNAs (28%) (Sup. Figure 4). More details concerning gene expressions and tissue specificity are available in Sup. Table 8 and 9 for the human and the chicken respectively. On the 1,555 putative orthologous lncRNAs considered as one-to-one between the human and the chicken using the method 2 (Table 1), 1,159 human lncRNA identifiers were found in the GTEx data (Sup. Table 10). Of this set, 39 lncRNA pairs were unexpressed in the 17 tissues for both species and were therefore removed from the analysis, bringing the number of analyzable lncRNA to 1,120 pairs. As shown in the Table 2 and Figure 4B-*i*, the expression values of each orthologous putative lncRNA constituting the pair were compared for the 17 tissues (see Sup. Table 11 for more details). On average, out of the 1,120 pairs considered for each tissue, 46% pairs are unexpressed in the 17 tissues for both species. Among the remaining 606 (54%) pairs expressed in at least one species, an average of 163 (25%) pairs were expressed in both species and 94 (15%) were in the same expression category (“Weak Expression” with expr. ∈ ]0.1, 1] TPM or “Standard expression” for expr > 1 TPM). Interestingly, 1,031 (92%) of the pairs were found as expressed in the testis for at least one species and 492 (48%) in both species, the only tissue displaying such a value. In comparison, a control set of 1,111 pairs of presumed non-orthologous lncRNAs was established, and on average, 565 (51%) were expressed in at least one species with a total of 82 (13%) being expressed in both. A significant enrichment (t-test, 25% *vs* 13%, p-value < 10^-5^) of lncRNA expressed in both species is consequently observed for lncRNA pairs assumed to be orthologous. Even if lncRNA and PCG are known to display different patterns of expression, two sets of 1,149 and 1,151 PCGs, including orthologous pairs and random pairs respectively, were analyzed for comparison with lncRNA. On average, 1,080 (94%) and 1,085 (94%) of these pairs appeared to be expressed in at least one species but 970 (90%) and 623 (57%) were expressed in both species respectively. Therefore, the significant order of magnitude (around a factor 2) between orthologous and non-orthologous PCG pairs (t-test, 90% vs 57%, p-value < 10^-17^), is near to the one observed for lncRNA pairs (25% *vs* 13%). Note that if the expression distribution observed between tissues tend to be similar for all the tissues, the testis shows a different pattern where the number of genes expressed is higher than for other tissues.

**Table 2.**
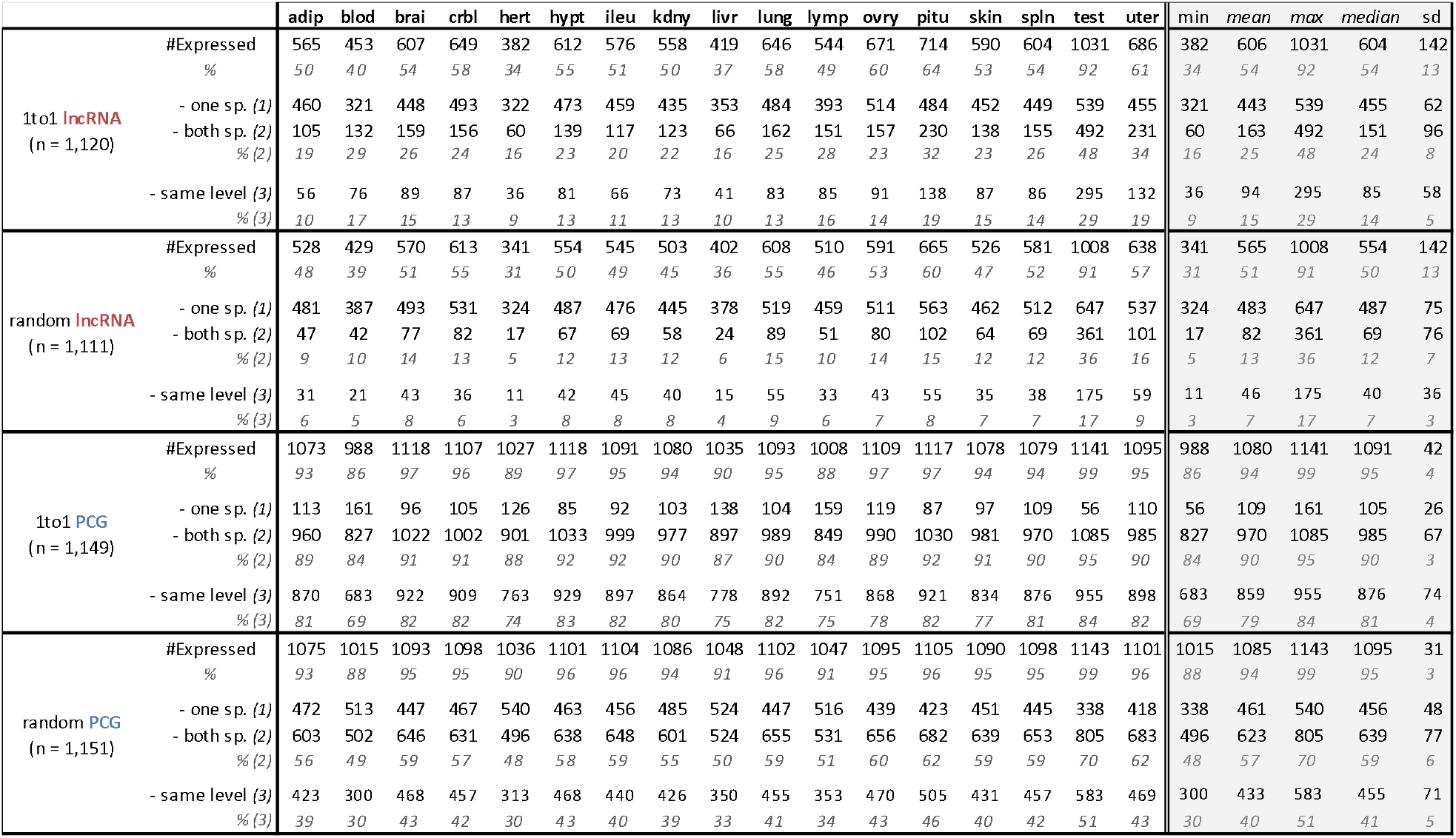
Number of lncRNAs or PCGs, orthologous or randomly selected, expressed between the chicken and the human for the 17 tissues studied. “#Expressed”: genes expressed in the given tissue for at least one specie. sp.: species. “same level”: genes with expressions belonging to the same class, “Weak” or “Standard” expression. Tissues abbreviations are available in the Sup. Table 7.

To observe the expression pattern across the 17 tissues, the co-expression between the 1,159 putative orthologous lncRNA pairs were computed (Sup. Table 10). In fact, a total of 798 pairs were analyzed, with each of the two lncRNAs showing expression in at least one tissue for each species. As show in the Figure 4B-*ii*, 151 (18.9%) and 37 (8.7%) pairs had a co-expression ≥ 0.5 and 0.7 respectively. To compare, sets of 1,159 non-orthologous lncRNAs and PCGs and a set of 1,159 orthologous PCG have been studied: 76 (6.6%) and 25 (2.2%), 76 (6.6%) and 12 (1.0%) and 522 (45.0%) and 185 (16.0%) pairs had a co-expression ≥ 0.5 and 0.7 respectively. Finally, since expression pattern can provide some information on a potential biological role of the lncRNA, the tissue-specificity was studied. Among the 798 pairs, 205 (25.7%) had at least one shared tissue among the most highly expressed tissue groups (*i.e.*, first break) but more than 79% were observed for the testis. Note that 136 pairs were detected as having a ≥ 0.90 in both species.

To illustrate the interest of gene expression conservation between the human and chicken (Figure 4B-*iii*), the human lncRNA (ENSG00000225386), named AC099754.1 in the GTEx version (GRCh38.p10) but identified as LRRC3B-AS1 in the latest version of the human genome (GRCh38.p14), and its putative chicken ortholog (LOC112531717) were used. This pair of putative orthologous lncRNAs presents a significant co-expression across tissues ( = 0.71, p-val < 0.05). Interestingly, while the gene is considered TS in humans (τ = 0.92), it is not in chickens (τ = 0.87). However, using the notion of "break" presented in Materials and Methods, both lncRNAs show 4 shared tissues in their first expression group associated with the cerebral system, including cerebellum (crbl), hypothalamus (hypt), brain (brai) and pituitary gland. This "first break group" is thus clearly distinct from other tissues, with a fold change of 13.44 in chickens when the median of each group is compared, and an absence of expression in other tissues in humans. These lncRNAs are both associated to a conserved PCG in divergent/antisense configuration: LRRC3B, showing the same expression pattern as the neighboring lncRNA and known to be related to brain function such as cholesterol metabolisms [20] and Down Syndrome [21, 22]. On the other hand, OTX2-AS and its putative homologous in chicken show a significant co-expression across tissues around 0.56 (p-val < 0.05). Both are considered as tissue-specific with a τ = 0.96 and τ = 0.95 in the chicken and the human respectively for the hypothalamus, what is consistent with OTX2 known functions (Figure 4B-*iv*).

## DISCUSSION

By combining three approaches—two based on synteny and one on multiple alignment—lists of potentially conserved lncRNAs were established, either between two species or among several phylogenetically distant species. A major limitation throughout this study is the variability in genome assemblies and associated annotations. They both can evolve rapidly, leading to changes in the established lists of putative orthologous lncRNAs [23].

Regarding genome assemblies, the periodicity of their updating could tend to diminish in the years to come, as new sequencing technologies lead to the creation of increasingly precise and complete assemblies, such as T2T assemblies, as demonstrated by the human assembly published in 2022 [24], or that of the chicken for example [25]. In addition, projects such as the Earth BioGenome Project [26] and the Darwin project [27], which consist in automatically sequence a large number of species, aim to provide reference genomes for a large panel of species. In terms of genome annotation, the democratization of long-read technologies [28] on RNA molecules and capture techniques for RNA lowly expressed [29, 30], as lncRNA, should also improve their accuracy over time, by refining gene models, in particular for PCGs, and enabling the annotation of previously unidentified genes such as for lncRNAs. However, once an assembly is selected as the reference, discrepancies are observed between the gene models provided by the reference databases such as Ensembl-EMBL and NCBI-Refseq. Specifically, protein-coding gene (PCG) models primarily differ at the transcript model level, whereas long non-coding RNA (lncRNA) gene models exhibit differences at both the transcript and gene loci levels. This observation underscores the interest of unifying the various annotations, as exemplified by the enriched chicken genome annotation used [31]. It should be noted that, for a given reference database, even if different versions of the genome annotation are available, only slight changes are observable independently of gene biotypes [23, 31]. These limitations concerning genome assemblies and annotation have consequently a technical impact on the three methods implemented in this study.

The one relying on the two-neighboring conserved PCGs (referred as “method 1“) is effective in identifying well-conserved lncRNAs, although it can be stringent. In phylogenetically distant species, the number of conserved protein-coding genes (PCGs) is often low, making it challenging to identify syntenic lncRNAs. Additionally, because algorithmically the lncRNA is used firstly to find the surrounding orthologous PCGs, they might not be conserved, leading to the exclusion of the lncRNA from the analysis. An alternative approach would involve the removal of all non-orthologous protein-coding genes (PCGs), thereby utilizing only 1-to-1 orthologous PCGs between species of interest, followed by the examination of conserved long non-coding RNAs (lncRNAs). However, this approach would inherently result in a reduction of PCGs used as anchors and an increase in the size of the delimited genomic areas. Consequently, this would favor many-to-many orthology relationships for the lncRNAs, which necessitate manual analysis due to their complexity. This effect would be further amplified when a larger number of species or species with significant phylogenetic divergences are considered. Moreover, in a subsequent step, the relative genomic orientation of the PCG-lncRNA-PCG triplet is taken into account to reinforce the putative orthologous relationship established. However, genomic rearrangements can alter the relative orientation, particularly when the distance between genes is substantial. Although it is possible to relax the gene orientation criteria, doing so is challenging with a large number of species and results in a reduction in precision.

To address the constraints associated with using two orthologous PCGs (“method 1”), the second approach (“method 2”) involves examining the nearest conserved PCGs and their configuration with the lncRNA of interest. This method is also influenced by annotation quality and the presence of gene models but the anchoring of a single PCG limits this impact. However, the classification, following FEELnc standards [32], is nevertheless dependent of the quality of gene models and especially associated isoforms. For instance, the classification requires distinguishing between antisense and convergent transcripts or between genic and intergenic ones, despite often poorly defined UTRs [33]. Relaxing this classification, as performed in this study, introduces user-dependent variability, which may affect the results based on user-imposed restrictions, however, the application of a strident configuration can also make relationships all the more certain.

Finally, the last approach (“method 3”), based on alignment, is less dependent on annotations but still relies on identified lncRNAs in the source species to find orthologous genomic regions through multiple sequence alignments (MSA). Constructing an MSA for numerous species, especially distant ones, is resource-intensive [34]. As an alternative, MSAs provided by reference resources such as Ensembl can be used, though not all species may be included, and managing the required assembly and annotation versions can be challenging. Moreover, the conservation of a genomic sequence does not guarantee the presence of the lncRNA in the target species.

Overall, while identifying putative orthologous relationships using each method is a preliminary step, finding this relationship through several or all methods provides stronger support. In this study, lists of potentially orthologous lncRNAs have been established and, as observed in the literature [4, 35, 36], the number of orthologous lncRNAs appears higher for well-annotated and phylogenetically close species, and conversely, more diffuse for species that diverged earlier. However, assuming an orthologous relationship does not necessarily imply that the lncRNAs or their functions are conserved, necessitating further confirmation. Nevertheless, if putative orthology is found in several species, particularly phylogenetically distant ones, the reliability of the assumed relationship is increased as presented with the OTX2-AS1 example.

To further characterize putative lncRNA orthologous pairs, additional functional analyses can be performed. Two main propositions are presented in the manuscript: *i)* the analysis of sequence conservation by functional patches, and the ii) conservation of gene expression profiles. For the latter, assuming that genomic proximity (i.e., between a lncRNA and a PCG) and significant expression correlation indicate common regulation or involvement of the lncRNA in regulating the PCG, this mechanism may be potentially conserved through evolution [37].

Regarding the analysis of conserved sequences *(i)*, the lncLOOM tool [8] is effective, although the precision of the gene models studied is crucial. The search for conserved motifs is additive, after reclassification of sequences according to their divergence or not. If a motif is not found in one step, it will not be searched for in the subsequent steps. The choice of the reference sequence on which the order is established is of prime importance as mentioned in Ross et al. [8], and as demonstrated here by the divergence of results observed for OTX2-AS1 depending on whether the human or chicken sequence is used as the reference. An alternative would be to test all the potentially orthologous sequences as a reference and observe all the motifs conserved between species. However, the imprecision of certain sequences could introduce bias, and restricting to sequences from the best-annotated species, such as human, mouse, or zebrafish, or from the species of interest, such as the chicken in this case, appears to be a better choice. Thus, while the conservation of small functional patches can provide additional support for the orthology relationship between potentially conserved lncRNA sequences, it can also provide functional information. Conserved zones can be associated with binding zones for regulatory elements, like miRNA as available via lncLOOM/Targetscan or RBP-binding sites as presented in Huang et al., 2024 [14]. Functional enrichment analyses can then be performed for example via the miRNA Enrichment Analysis Annotation Tool (miEAA [38]). Moreover, under certain conditions, it is possible to observe whether the expression profiles of the regulatory elements are consistent with those of the genes under study.

For the observation of gene expression profiles *(ii)*, this first requires a dataset for the two (or more) species of interest containing both a fairly substantial number of tissues that can be considered equivalent/proximal, in order to observe expression profiles and address the notions of tissue-specificity. Furthermore, while it is now recognized that lncRNAs are much more tissue-specific than PCGs [1, 31, 39], such tissues must be present in the set of tissues under study, otherwise a significant number of lncRNAs will be discarded. Considering these different limitations, a first level of analysis is to determine whether the two lncRNAs considered as orthologs are expressed in the same tissues. A further nuance can be added by comparing their expression thresholds, specifically by observing whether they are weakly expressed (with a value of 0.1 to 1 TPM typically used) or more highly expressed (> 1 TPM). Although it is recognized that expression quantification may be biased according to the sample panel, the RNA sequencing techniques employed, or the number of reads sequenced, this initial approach helps to support the reliability of the orthologous relationships with two times more pairs expressed in the same tissues than observed in the control set of non-orthologous pairs. In a second step, the expression of the lncRNA pair can be observed across the whole tissue**s**, benefiting from great variability of expression across tissues, and the rank correlation can be calculated to observe whether lncRNAs share common profiles. Three times more orthologous lncRNA pairs are observed as co-expressed (>0.5) compared to the control set of non-orthologous pairs. In-depth study of these profiles can also identify pairs with common tissue-specificity, providing a first indication of function, as illustrated by the expression profile analysis of causal genes associated with Mendelian traits [31]. However, tissue specificity is a relative measure, dependent on multiple factors, especially the metric employed [40]. Indeed, some of them provide a score and a single tissue, omitting tissues with shared/common functions. The approach described in the manuscript, which consists in observing tissues that show a break in comparison to others, partly addresses this problem.

Finally, the work presented here provides a variety of three approaches for inferring orthologous relationships between lncRNAs potentially conserved at different phylogenetic scales. Additional analyses based on functional short-motif enrichment across the 13 species using the LncLOOM tool and co-expression patterns through 17 shared tissues between human and chicken provide some

information related to functionality of these orthologous lncRNAs. However, molecular biology experiments are needed to further investigate our understanding of the function of these orthologous lncRNAs.

## METHODS

### Assemblies and genomic annotations

Genomic data included annotated assemblies from 13 species of diverse taxa. Species selected included *i)* model organisms such as *Homo sapiens* (human), *Mus musculus* (mouse), and *Danio rerio* (zebrafish) and *ii)* domestic species with two clades represented: *a) Boreoeutherian* with *Canis lupus familiaris* (dog), *Bos taurus* (cow), *Equus caballus* (horse), *Sus scrofa* (pig)*, Capra hircus* (goat), and *b)* Aves with *Gallus gallus* (chicken), *Taeniopygia guttata* (zebra finch), *Meleagris gallopavo* (turkey), *Struthio camelus australis* (ostrich), and *Anas platyrhynchos* (duck). For each species, the genome assembly and corresponding annotation available in Ensembl version 109 were used, except for *Gallus gallus*, where a custom lncRNA-enriched annotation published by Degalez et al. in 2024 [31] was employed. Briefly, this annotation integrates data from six different resources, including the two reference annotation databases EMBL-EBI/Ensembl (v107 equivalent to the v109) and NCBI/RefSeq (v106), to provide a comprehensive atlas of lncRNA genes. Additional details regarding the specific assembly and annotation versions and availability for the 13 species are provided in Sup. Table 12.

### Identification of one-to-one orthologous PCGs

Gene orthology was extracted by species pair using BioMart [41] from Ensembl (v109). Since the vast majority of orthology links provided are for PCG, only these have been conserved.

### Quantification of one-to-one orthologous PCGs proportion between two species

To quantify similarity between the set of one-to-one PCG between every pair of species, the Jaccard-index was calculated as follow:

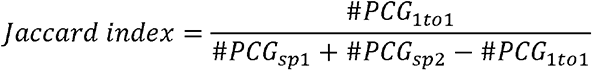

with #*PCG_spi_* the numbers of PCGs in species *i* and #*PCG_1to1_* the number of one-to-one PCGs between both species. By definition, a Jaccard-index value close to 1 indicates high similarity or overlap between the sets, while a value closer to 0 suggests minimal overlap.

### Analysis of N-to-N orthologous lncRNAs according to their number between both species

The systematic classification of orthology relationships between lncRNAs across species was inspired by the approach typically used for orthologous PCGs. Based on the number of lncRNAs identified in the source and target species, a categorical label was assigned (Figure 2D). The possible categories include *i)* “one-to-zero” or “many-to-zero” when a single or several lncRNA(s) in one species respectively have no orthologous lncRNA in the other species; *ii)* “one-to-one” when a single lncRNA is observed in both species; *iii)* “one-to-many” when a single lncRNA in one species are linked to multiple lncRNAs in the other species; *iv)* “many-to-one” for the opposite case and *v)* many-to-many for multiple lncRNAs in both species. For the many-to-many case, an additional tag was implemented to indicate if the number of lncRNAs was equal or different between species.

### FEELnc orientation analysis of lncRNAs according to nearest PCG

LncRNA transcripts were first classified relatively to their closest PCG transcript using the “FEELnc_classifier” function of FEELnc v.0.2.1 [32, 42] with a maximum window of 100 kb (default setting). The classification for gene models was performed by combining the transcript results and the “tpLevel2gnLevelClassfication” function from FEELnc. However, due to differences in genome annotation between species and since lncRNA transcripts are sometimes poorly known, *e.g.*, an antisense lncRNA-PCG pair in one species being considered divergent in another, the FEELnc configuration nomenclature was relaxed. Therefore, as shown in Sup. Figure 2, four categories were defined: *i)* “Conv” for tail-to-tail and *ii)* “Divg” for head-to-head oriented gene and *iii)* “SS.up” and *iv)* “SS.down” for upstream and downstream lncRNAs respectively, in the same orientation as the PCG. Each category is then divided in two groups according to the distance between the lncRNA and PCG TSSs. If the distance is less than 5,000 bp or if the lncRNA TSS is within the PCG boundaries, it is considered as genic (“lncg”) otherwise intergenic (“linc”).

### Methods to infer lncRNA orthology relation

#### Method 1: “Orthology-by-synteny” using the two orthologous flanking PCGs (*Figure 2A*)

For each species, lncRNAs are associated to the nearest upstream and downstream PCG. Then, for each pair of species, lncRNAs located between pairs of PCGs with one-to-one orthology relationships were extracted. If the relative orientation and strand configurations of the lncRNAs and their flanking PCGs is concordant, *i.e*, identical or in reverse, lncRNAs of both species are considered as putative positional orthologs. The type of relation is established as presented in the “N-to-N orthologous lncRNA analysis” previous part.

#### Method 2: “Orthology-by-synteny” using one orthologous flanking PCGs and the conserved FEELnc orientation of the lncRNA-PCG pair (*Figure 2B*)

For each species, lncRNAs were classified relatively to the nearest PCG and according to the relaxed FEELnc nomenclature (“FEELnc orientation analysis of lncRNAs according to nearest PCG” paragraph) presented previously. LncRNAs associated to PCGs with one-to-one orthology relationships and with the same FEELnc orientation were then considered as putative orthologs. The type of relation is established as presented in the “N-to-N orthologous lncRNA analysis” previous part.

#### Method 3: “Orthology-by-genome alignment” using Ensembl/Compara and the Mercator PECAN alignment of amniotes group (*Figure 2C*)

The Perl API provided by Ensembl was used to easily access to the Compara database [43]. For each species, the coordinates of all lncRNAs were extracted. After loading the registry associated to the last Ensembl version (here v111), adaptors associated to the species concerned were used. Genomic alignment blocks were retrieved using the “PECAN” method and for the “amniotes” group. Briefly, the Mercator-Pecan algorithm [16] include two steps: 1) the orthology map between the genomes of interest is defined using Mercator which uses orthologous coding exons to define blocks of orthologous segments; 2) Orthologous blocks are then aligned using Pecan. The genomic coordinates of all lncRNAs provided as input were used to fetch corresponding genomic alignment blocks, which were then further processed to obtain relevant genomic alignment information for all species included in the 63 amniotes group and for which a match has been observed. For this study and using Ensembl v111, 10 species of interest were available (*Canis lupus familiaris, Equus caballus, Capra hircus, Sus scrofa, Mus musculus, Homo sapiens, Taeniopygia guttata, Anas platynrhynchos, Meleagris gallopavo, Gallus Gallus*) excluding *Bos taurus*, *Struthio camelus australis* and *Danio rerio* from the analysis. By construction, the result of this multi-species alignment is a consensus sequence based on the 63 amniotes, but only the block alignment for our 10 species of interest is displayed here.

### Conservation of short motifs (k-mers) between species sequences

For lncRNA of interest, LncLOOM v2.0 [8, 44] was used to identify combinations of short motifs that are conserved across different species of interest. FASTA sequences for lncRNAs exons identified as one-to-one putative orthologs by method 1 or method 2 was used as input. If the lncRNAs had several transcripts, all the exonic parts were considered. The “--targetscan” option was applied to annotate motifs corresponding to conserved miRNA sites according to TargetScan [13]. The k-mers search was performed for motif sizes ranging from 20 (--startw 20) to 6 (default: --stopw 6). The enrichment analysis for the miRNA binding sites detected by TargetScan was conducted using the miRNA Enrichment Analysis Annotation Tool (miEAA) [38].

### Expression data among 17 tissues (limited to chicken and human comparison)

Expression analyses were conducted exclusively between chicken and human. For human data, expression in TPM were derived from the RNA-seq expressions provided by the human GTEx Analysis v8 project [45, 46]. Since the gene annotation used for the human in this study (GRCh38.p13) and the one used by GTEx (GRCh38.p10) are different, only genes with the same gene identifiers in both annotations have been kept (n = 55,525 genes; 19,214 PCGs and 13,634 lncRNAs). Chicken expression in TPM were obtained from the chicken atlas published by Degalez et al., 2024 [31] and accessible on the GEGA user-friendly online tool (https://gega.sigenae.org/) [47]. For both *i)* inter-tissue and *ii)* intra-tissue analysis, a list of 17 tissues with at least 8 samples per tissue for each species and common to both species was established (see Sup. Table 6). A gene was considered as expressed in a tissue if its expression was greater than 0.1 TPM for at least 80% of the samples.

### Co-expression indicator

All co-expression analyses were conducted using expression in TPM and the Spearman correlation ( ). Genes were considered as co-expressed for a | | ≥ 0.7 after that p-values were corrected for multiple testing using the Benjamini-Hochberg method and applying a false discovery rate of 0.05.

### Tissue-specificity analysis (limited to chicken and human)

Multiple criteria based on the tau and FC metrics were used to assess the tissue-specificity of each gene:

*i)* the tau metrics [48], assessed using raw (*τ*) or log10 median tissue expression (*τ*_log10_) in TPM are defined as follows:

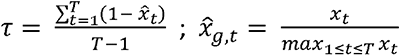

with the *x_t_* raw or log10 median expression of the gene of interest in the tissue *t* and among the *T* tissues.

A gene was considered as tissue specific for or *τ* or *τ*_log10_ or ≥ 0.90;

*ii)* the FC between the first and second most expressed tissues;

*iii)* it should be noted that a gene may show relatively similar expression between some tissues, often functionally close, while also showing significant expression differences with other tissues. We therefore introduced the notion of breaks, defined here as a difference in expression by a factor of 2 (*i.e.*, FC ≥ 2) between two tissues, after ranking the expression of the tissues in decreasing order. We then introduced a third criteria to consider these breaks: the FC between the two first groups separated by a break, which was calculated though three different ways, considering: *a)* the least expressed tissue in the first group and the most expressed tissue in the second one; *b)* the most expressed tissue in both groups; *c)* the median expression for both groups. Tissue-specificity results for all the genes are available in Sup. Table8-9. These different TS criteria can be found in the GEGA user-friendly tool [47] navigating across 47 tissues expressions in chicken.

## DATA AVAILABILITY

Expression data and the custom chicken gene-enriched annotation underlined in this article are available via the GEGA website at https://gega.sigenae.org/ and through the “Download” tab. Scripts used for the identification of putative orthologous lncRNA between species are available on Github at https://github.com/fdegalez/lncrna_PutOrtho.

## ACKNOWLEDGMENTS

We would like to thank OpenMoji community which provides an open source and independent emoji system that we used for the figure design.

## AUTHOR CONTRIBUTIONS

FD and SL conceived and coordinated the study. SL acquired funding for this research. FD and SL carried out the whole bioinformatics analysis. CA and LL carried out the wet analysis. FL was responsible for the computational infrastructure. FD and SL drafted the manuscript and figures. CA, LL, FL helped to improve the manuscript. All authors reviewed and approved the final version.

---

**Sup. Figure 1.**
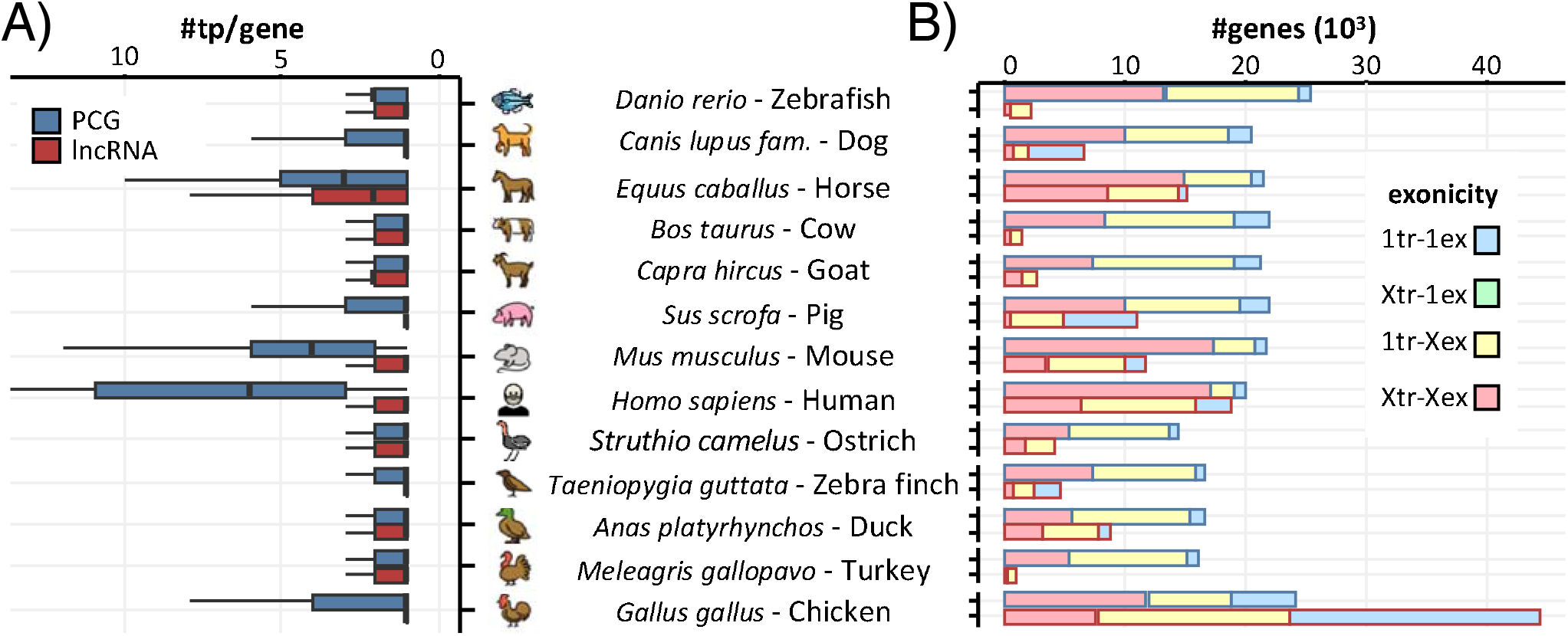
PCGs and lncRNA characteristics of the 13 species. (a) Number of transcripts associated to each PCGs (blue) and lncRNAs (red). (b) Number of PCGs and lncRNA supported by one (“1tr”) or more (“Xtr”) transcripts and with one (“1ex”) or more (“Xex”) exons.

**Sup. Figure 2.**
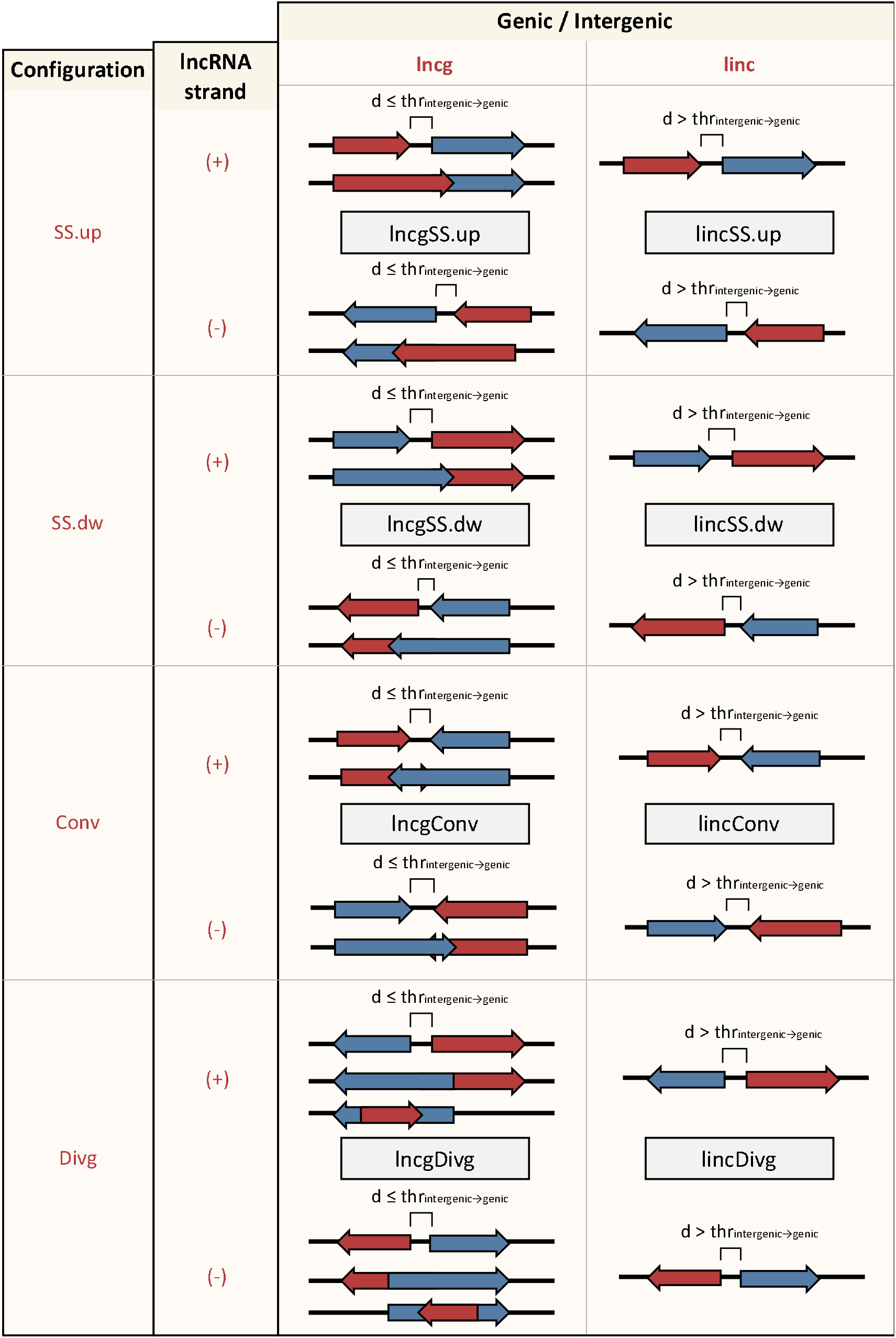
Classification for lncRNA according to the nearest PCG. LncRNAs and PCGs are represented in red and blue respectively. The arrow orientation corresponds to the gene strand. d: minimum distance between the lncRNA and PCG. thr_intergenic→genic_: threshold distance to distinguish genic and intergenic gene.

**Sup. Figure 3.**
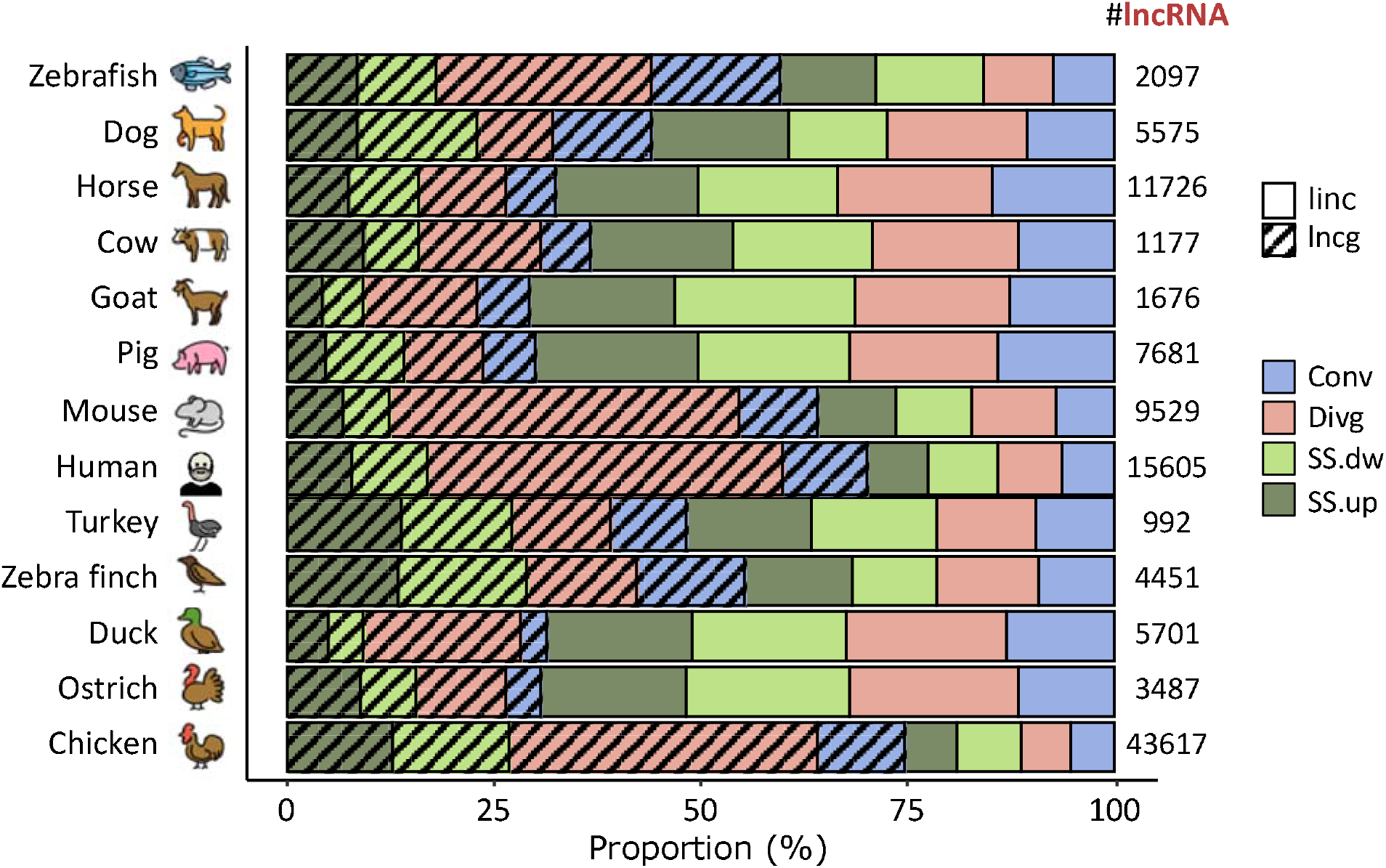
Proportion of lncRNA classified following the FEELnc relaxed classification for the 13 species. #lncRNA: total number of lncRNA classified by FEELnc. linc: intergenic; lncg: genic. Conv: convergent; Divg: divergent; SS.dw: same-strand downstream; SS.up: same-strand upstream.

**Sup. Figure 4.**
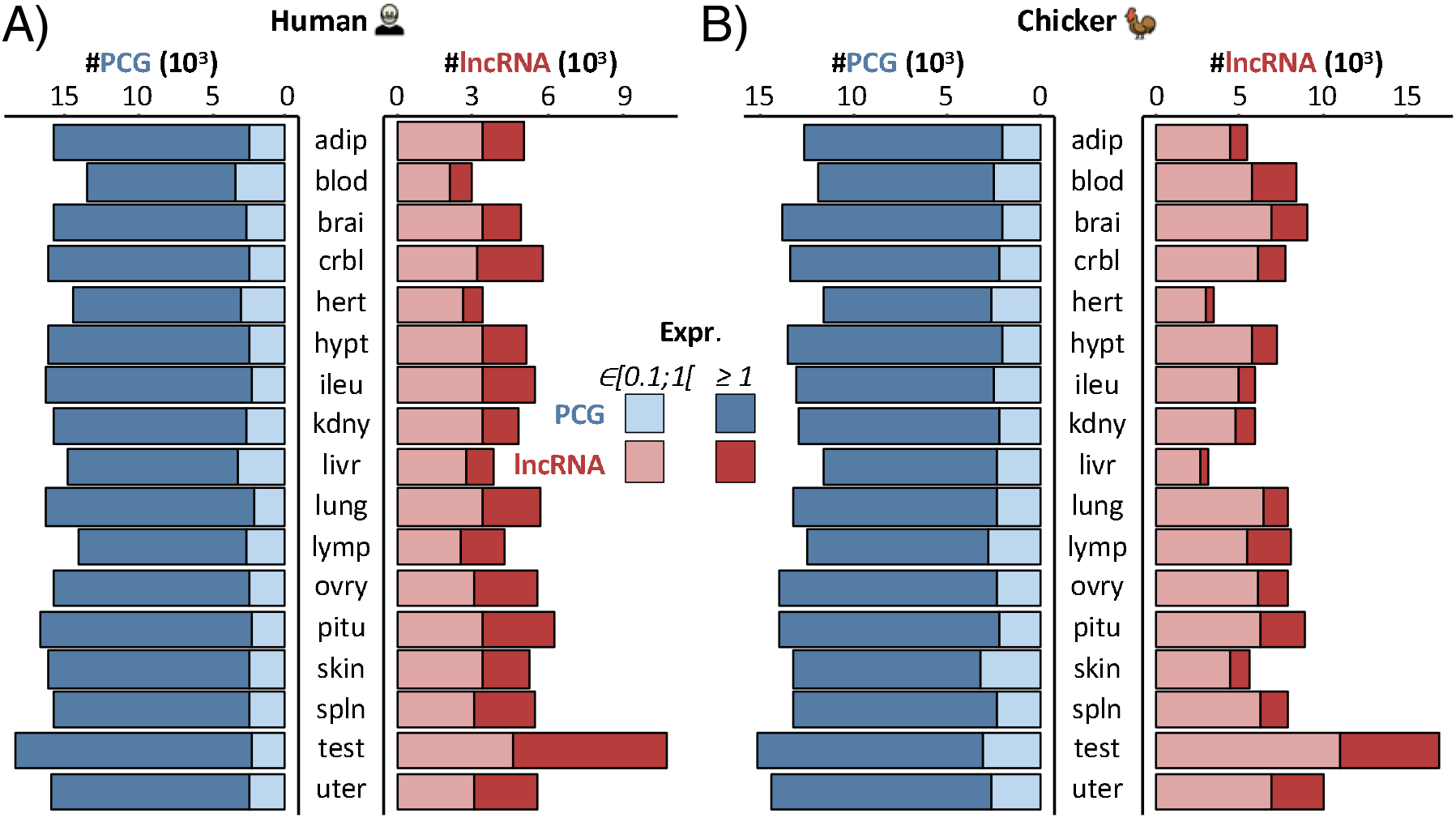
Gene expression through the 17 equivalent tissues for the human and the chicken. Number of PCGs (blue) and lncRNAs (red) with an expression in TPM∈ [0.1;1[ (light shade) or ≥ 1 (dark shade) for the 17 tissues and for the human (a) and the chicken (b) respectively.

**Sup. Table 1** – Number of lncRNAs surrounded by 0, 1 or 2 PCG orthologs when analyzed by species pair and method 1.

**Sup. Table 2** – Number of lncRNA orthologous relations found by each method for all the 13 species. For method 1 and method 2, “1to1” refers to lncRNA with a “one-to-one” ortholog and “many” for lncRNA with a “many-to-one”, “one-to-many” or “many-to-many” relation. Union (∪) and intersection (⋂) for method 1 and 2 (1-2) and all methods (1-2-3). NA: Not Applicable.

**Sup. Table 3** – For each species, number of lncRNAs in each FEELnc category as defined in the method 2.

**Sup. Table 4** –Additional information for OTX2-AS1 identification in the 13 species and using the three methods presented.

**Sup. Table 5** - N–ucleotide motifs and miRNAs identified and their respective depths for the OTX2-AS1 lncRNA analyzed by LncLOOM, considering the human and chicken sequences as reference respectively.

**Sup. Table 6** – Functional enrichment through miEAA for miRNAs identified for OTX2-AS1 via TargetScan et LncLOOM.

**Sup. Table 7** - Inf–ormation on the 17 tissues considered as equivalent between the human and the chicken.

**Sup. Table 8** – Expression and tissue-specificity indicators computed for the 17 equivalent tissues for the human. Tissues abbreviations are available in the Sup. Table 7.

**Sup. Table 9** – Expression and tissue-specificity indicators computed for the 17 equivalent tissues for the chicken. Tissues abbreviations are available in the Sup. Table 7.

**Sup. Table 10** – Expression, tissue-specificity and co-expression indicators computed for the 17 equivalent tissues for lncRNA putative orthologous pairs between the human and the chicken. Tissues abbreviations are available in the Sup. Table 7.

**Sup. Table 11** – Number of PCGs, lncRNAs and 1to1 putative orthologous lncRNAs according to the three expression categories (TPM < 0.1; ∈ [0.1;1[ or ≥ 1) and through the 17 equivalent tissues for the human and the chicken. Tissues abbreviations are available in the Sup. Table 7.

**Sup. Table 12** – Information on the annotation and assembly of the genomes used. Download links from reference databases.

## REFERENCES

1. Derrien T, Johnson R, Bussotti G, Tanzer A, Djebali S, Tilgner H, et al. The GENCODE v7 catalog of human long noncoding RNAs: Analysis of their gene structure, evolution, and expression. Genome Res. 2012;22:1775–89.

2. Uszczynska-Ratajczak B, Lagarde J, Frankish A, Guigó R, Johnson R. Towards a complete map of the human long non-coding RNA transcriptome. Nat Rev Genet. 2018;19:535–48.

3. Statello L, Guo C-J, Chen L-L, Huarte M. Gene regulation by long non-coding RNAs and its biological functions. Nat Rev Mol Cell Biol. 2021;22:96–118.

4. Hezroni H, Koppstein D, Schwartz MG, Avrutin A, Bartel DP, Ulitsky I. Principles of Long Noncoding RNA Evolution Derived from Direct Comparison of Transcriptomes in 17 Species. Cell Reports. 2015;11:1110– 22.

5. Ulitsky I. Evolution to the rescue: using comparative genomics to understand long non-coding RNAs. Nat Rev Genet. 2016;17:601–14.

6. Quinn JJ, Chang HY. Unique features of long non-coding RNA biogenesis and function. Nat Rev Genet. 2016;17:47–62.

7. Rivas E. Evolutionary conservation of RNA sequence and structure. WIREs RNA. 2021;12:e1649.

8. Ross CJ, Rom A, Spinrad A, Gelbard-Solodkin D, Degani N, Ulitsky I. Uncovering deeply conserved motif combinations in rapidly evolving noncoding sequences. Genome Biology. 2021;22:29.

9. Sarropoulos ioannis, Marin R, Cardoso-Moreira M, Kaessmann H. Developmental dynamics of lncRNAs across mammalian organs and species. Nature. 2019;571:510–4.

10. Lagarrigue S, Lorthiois M, Degalez F, Gilot D, Derrien T. LncRNAs in domesticated animals: from dog to livestock species. Mamm Genome. 2022;33:248–70.

11. Muret K, Désert C, Lagoutte L, Boutin M, Gondret F, Zerjal T, et al. Long noncoding RNAs in lipid metabolism: literature review and conservation analysis across species. BMC Genomics. 2019;20:882.

12. Foissac S, Djebali S, Munyard K, Vialaneix N, Rau A, Muret K, et al. Multi-species annotation of transcriptome and chromatin structure in domesticated animals. BMC Biol. 2019;17:108.

13. Agarwal V, Bell GW, Nam J-W, Bartel DP. Predicting effective microRNA target sites in mammalian mRNAs. eLife. 2015;4:e05005.

14. Huang W, Xiong T, Zhao Y, Heng J, Han G, Wang P, et al. Computational prediction and experimental validation identify functionally conserved lncRNAs from zebrafish to human. Nat Genet. 2024;56:124–35.

15. EMBL EBI Ensembl/GENCODE. Multiple genome alignments. 2024. https://www.ensembl.org/info/genome/compara/multiple_genome_alignments.html. Accessed 28 Jun 2024.

16. Paten B, Herrero J, Beal K, Fitzgerald S, Birney E. Enredo and Pecan: Genome-wide mammalian consistency-based multiple alignment with paralogs. Genome Res. 2008;18:1814–28.

17. Romano GL, Platania CBM, Leggio GM, Torrisi SA, Giunta S, Salomone S, et al. Retinal biomarkers and pharmacological targets for Hermansky-Pudlak syndrome 7. Sci Rep. 2020;10:3972.

18. Qin N, Paisana E, Picard D, Leprivier G, Langini M, Custódia C, et al. The long non-coding RNA OTX2- AS1 promotes tumor growth and predicts response to BCL-2 inhibition in medulloblastoma. J Neurooncol. 2023;165:329–42.

19. Alfano G, Vitiello C, Caccioppoli C, Caramico T, Carola A, Szego MJ, et al. Natural antisense transcripts associated with genes involved in eye development. Human Molecular Genetics. 2005;14:913–23.

20. Shendre A, Wiener H, Irvin MR, Zhi D, Limdi NA, Overton ET, et al. Admixture Mapping of Subclinical Atherosclerosis and Subsequent Clinical Events Among African Americans in Two Large Cohort Studies. Circ Cardiovasc Genet. 2017;10:e001569.

21. Mills JD, Ward M, Kim WS, Halliday GM, Janitz M. Strand-specific RNA-sequencing analysis of multiple system atrophy brain transcriptome. Neuroscience. 2016;322:234–50.

22. Lockstone HE, Harris LW, Swatton JE, Wayland MT, Holland AJ, Bahn S. Gene expression profiling in the adult Down syndrome brain. Genomics. 2007;90:647–60.

23. Smith J, Alfieri JM, Anthony N, Arensburger P, Athrey GN, Balacco J, et al. Fourth Report on Chicken Genes and Chromosomes 2022. Cytogenet Genome Res. 2022;162:405–528.

24. Nurk S, Koren S, Rhie A, Rautiainen M, Bzikadze AV, Mikheenko A, et al. The complete sequence of a human genome. Science. 2022;376:44–53.

25. Huang Z, Xu Z, Bai H, Huang Y, Kang N, Ding X, et al. Evolutionary analysis of a complete chicken genome. Proceedings of the National Academy of Sciences. 2023;120:e2216641120.

26. Lewin HA, Robinson GE, Kress WJ, Baker WJ, Coddington J, Crandall KA, et al. Earth BioGenome Project: Sequencing life for the future of life. Proceedings of the National Academy of Sciences. 2018;115:4325–33.

27. The Darwin Tree of Life Project Consortium. Sequence locally, think globally: The Darwin Tree of Life Project. Proceedings of the National Academy of Sciences. 2022;119:e2115642118.

28. Marx V. Method of the year: long-read sequencing. Nat Methods. 2023;20:6–11.

29. Lagarde J, Uszczynska-Ratajczak B, Carbonell S, Pérez-Lluch S, Abad A, Davis C, et al. High-throughput annotation of full-length long noncoding RNAs with capture long-read sequencing. Nat Genet. 2017;49:1731–40.

30. Carbonell Sala S, Uszczyńska-Ratajczak B, Lagarde J, Johnson R, Guigó R. Annotation of Full-Length Long Noncoding RNAs with Capture Long-Read Sequencing (CLS). In: Cao H, editor. Functional Analysis of Long Non-Coding RNAs: Methods and Protocols. New York, NY: Springer US; 2021. p. 133–59.

31. Degalez F, Charles M, Foissac S, Zhou H, Guan D, Fang L, et al. Enriched atlas of lncRNA and protein- coding genes for the GRCg7b chicken assembly and its functional annotation across 47 tissues. Sci Rep. 2024;14:6588.

32. Wucher V, Legeai F, Hédan B, Rizk G, Lagoutte L, Leeb T, et al. FEELnc: a tool for long non-coding RNA annotation and its application to the dog transcriptome. Nucleic Acids Res. 2017;45:e57.

33. Amarasinghe SL, Su S, Dong X, Zappia L, Ritchie ME, Gouil Q. Opportunities and challenges in long- read sequencing data analysis. Genome Biol. 2020;21:30.

34. Chowdhury B, Garai G. A review on multiple sequence alignment from the perspective of genetic algorithm. Genomics. 2017;109:419–31.

35. Washietl S, Kellis M, Garber M. Evolutionary dynamics and tissue specificity of human long noncoding RNAs in six mammals. Genome Res. 2014;24:616–28.

36. Necsulea A, Soumillon M, Warnefors M, Liechti A, Daish T, Zeller U, et al. The evolution of lncRNA repertoires and expression patterns in tetrapods. Nature. 2014;505:635–40.

37. Engreitz JM, Haines JE, Perez EM, Munson G, Chen J, Kane M, et al. Local regulation of gene expression by lncRNA promoters, transcription and splicing. Nature. 2016;539:452–5.

38. Aparicio-Puerta E, Hirsch P, Schmartz GP, Kern F, Fehlmann T, Keller A. miEAA 2023: updates, new functional microRNA sets and improved enrichment visualizations. Nucleic Acids Research. 2023;51:W319–25.

39. Jehl F, Muret K, Bernard M, Boutin M, Lagoutte L, Désert C, et al. An integrative atlas of chicken long non-coding genes and their annotations across 25 tissues. Sci Rep. 2020;10:20457.

40. Kryuchkova-Mostacci N, Robinson-Rechavi M. A benchmark of gene expression tissue-specificity metrics. Briefings in Bioinformatics. 2017;18:205–14.

41. Smedley D, Haider S, Ballester B, Holland R, London D, Thorisson G, et al. BioMart – biological queries made easy. BMC Genomics. 2009;10:22.

42. Derrien T. tderrien/FEELnc. 2024.

43. Herrero J, Muffato M, Beal K, Fitzgerald S, Gordon L, Pignatelli M, et al. Ensembl comparative genomics resources. Database. 2016;2016:bav096.

44. lncLOOM. lncLOOM/lncLOOM. 2024.

45. GTEx Consortium. GTEx Portal- Bulk tissue expression. 2023. https://gtexportal.org/home/downloads/adult-gtex/bulk_tissue_expression. Accessed 4 Jan 2024.

46. THE GTEX CONSORTIUM. The GTEx Consortium atlas of genetic regulatory effects across human tissues. Science. 2020;369:1318–30.

47. Degalez F, Bardou P, Lagarrigue S. GEGA (Gallus Enriched Gene Annotation): an online tool providing genomics and functional information across 47 tissues for a chicken gene-enriched atlas gathering Ensembl & Refseq genome annotations. 2024;:2024.03.13.584813.

48. Yanai I, Benjamin H, Shmoish M, Chalifa-Caspi V, Shklar M, Ophir R, et al. Genome-wide midrange transcription profiles reveal expression level relationships in human tissue specification. Bioinformatics. 2005;21:650–9.

49. Kumar S, Suleski M, Craig JM, Kasprowicz AE, Sanderford M, Li M, et al. TimeTree 5: An Expanded Resource for Species Divergence Times. Mol Biol Evol. 2022;39:msac174.

