## Supplementary figures and images for "Cross-species orthology detection of long non-coding RNAs (lncRNA) through 13 species using genomic and functional annotations"

### Supplemental Figure 1

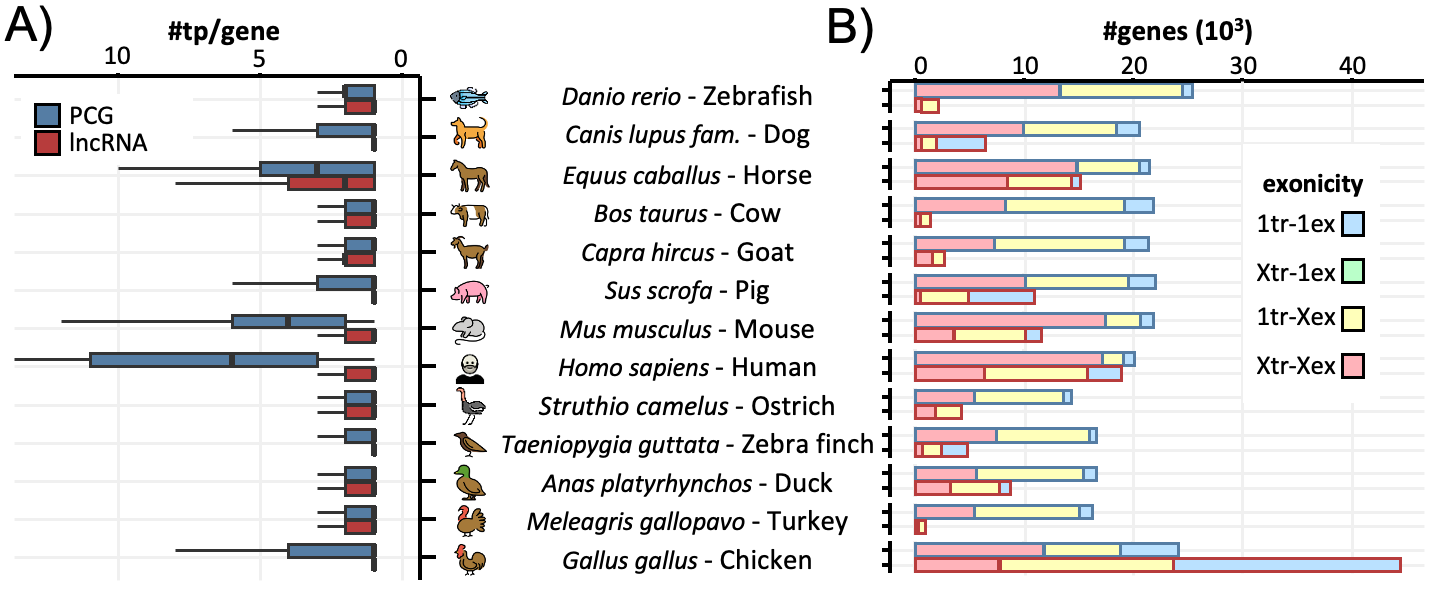

### Supplemental Figure 2

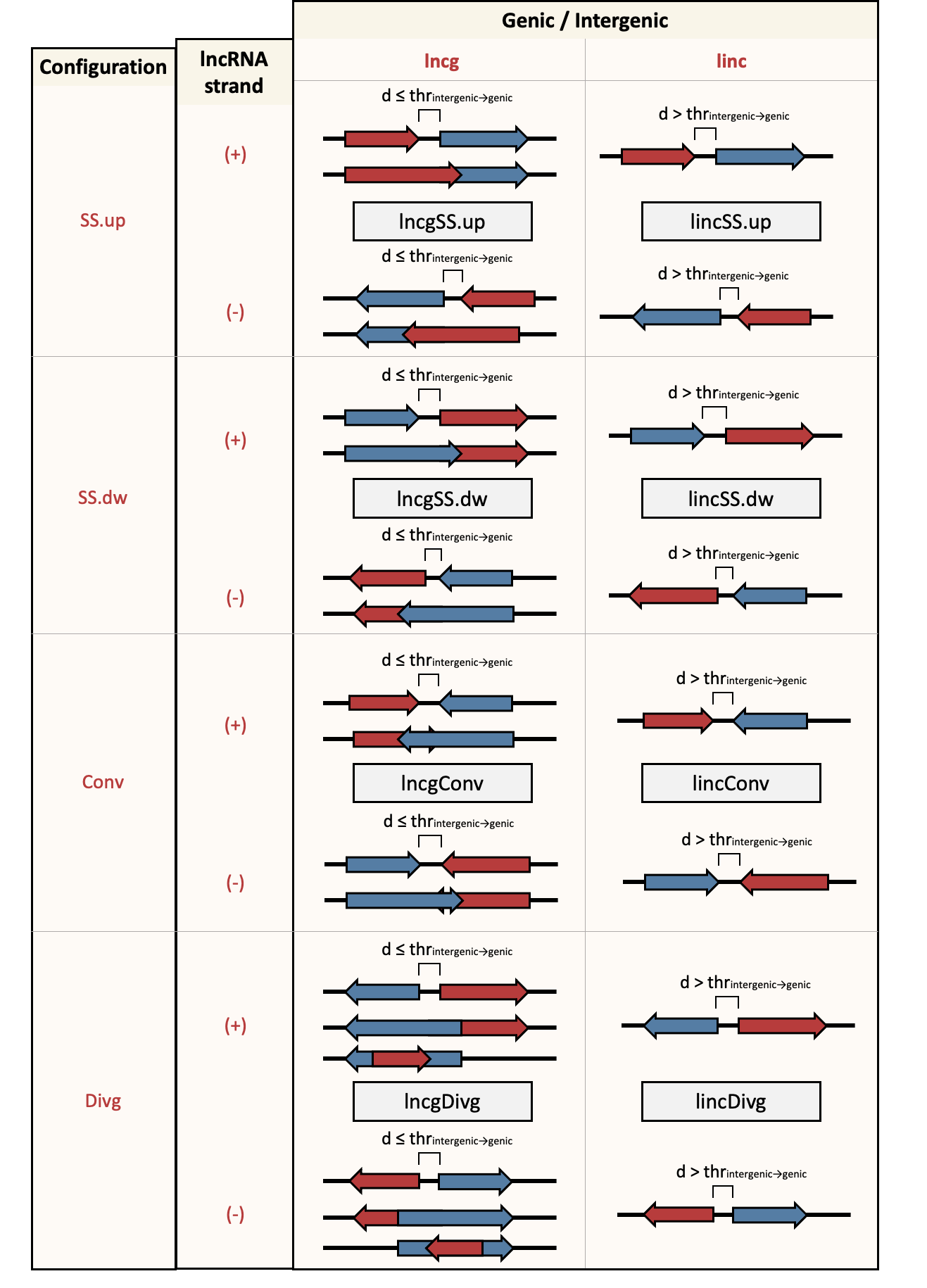

### Supplemental Figure 3

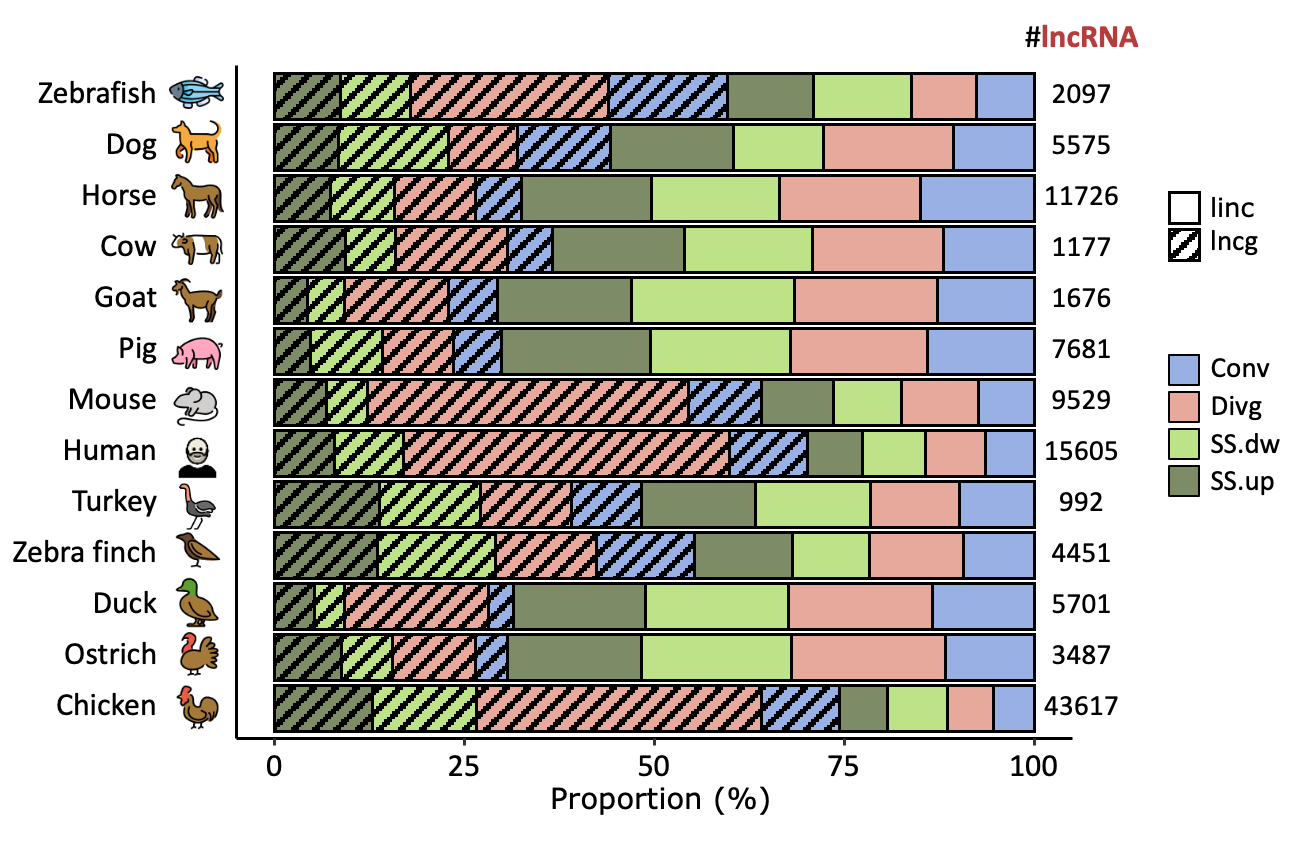

### Supplemental Figure 4

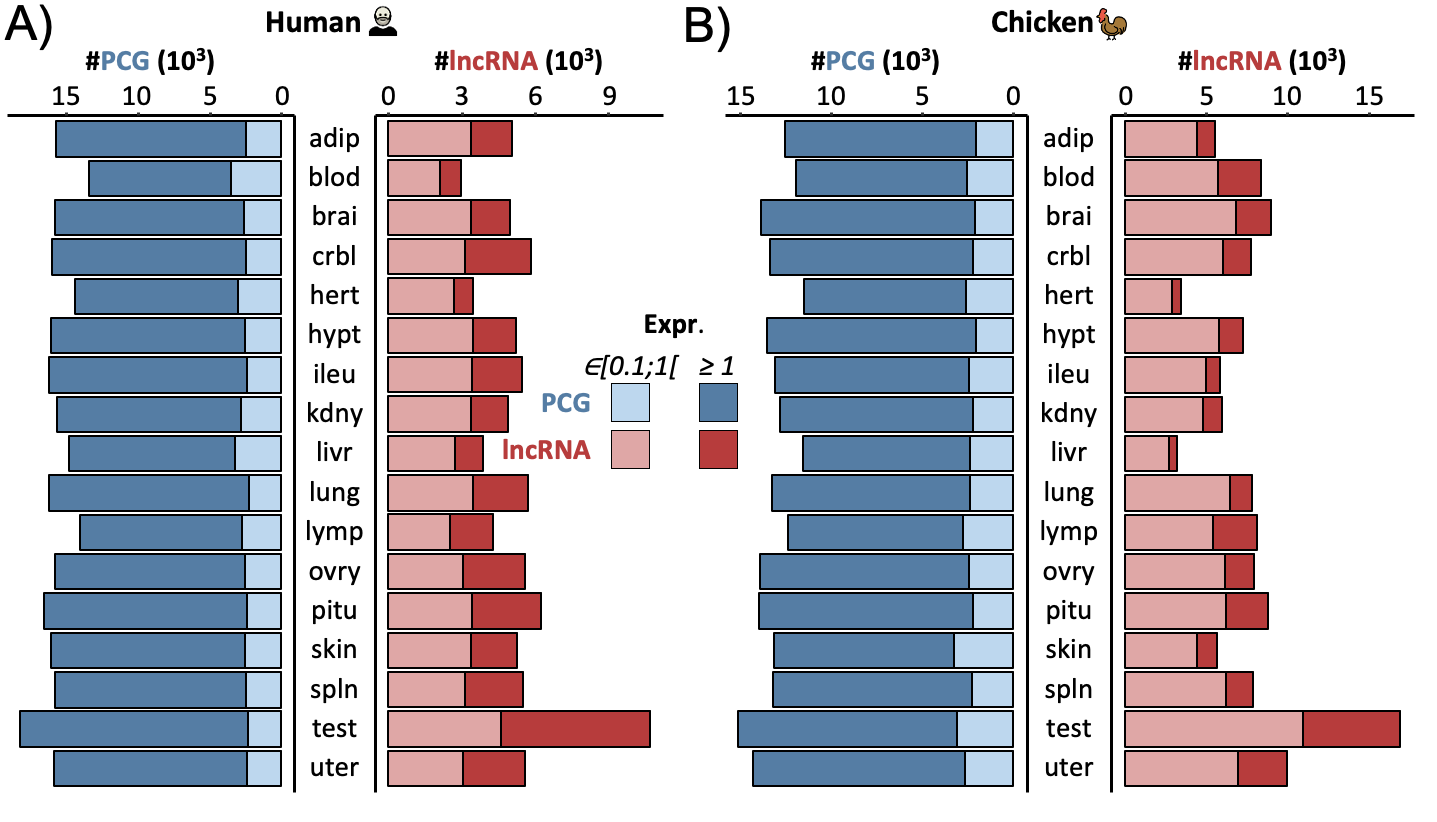
